# Hydroxy polyethylene glycol: a solution to evade human pre-existing anti-PEG antibodies for efficient delivery

**DOI:** 10.1101/2024.10.21.619346

**Authors:** Tianhao Ding, Jiaru Fu, Min Yang, Zui Zhang, Yinyu Ma, Ercan Wu, Zhiwei Guo, Shiqi Lin, Songli Wang, Xiaohua Liu, Bin Wang, Guanghui Li, Changyou Zhan

## Abstract

Polyethylene glycol (PEG) has been extensively utilized in food, cosmetics, and pharmaceutical fields, especially in the realm of nanomedicines, where it serves as a pivotal excipient for extending the nanoparticles circulation half-life. Contrary to its historical perception as non-immunogenic, pre-existing anti-PEG antibodies have been widely detected in human who even have never been exposed to PEGylated therapeutics, which considered to be associated with serious side effects of PEGylated nanomedicines including infusion reactions and other hypersensitive reactions. Herein, we elucidated the prevalence and distribution characteristics of pre-existing anti-PEG antibodies in 2074 human blood samples, and investigated its binding with PEG. Pre-existing anti-PEG antibodies were found to primarily recognize the PEG terminus, especially methoxy, which is the only PEG terminus contained in currently marketed PEGylated nanomedicines. While hydroxy PEG (OH-PEG) significantly evaded binding with pre-existing anti-PEG antibodies among most clinical samples. Noteworthily, substituting OH-PEG for MeO-PEG significantly mitigated complement activation of lipid nanoparticle (LNP) caused by pre-existing anti-PEG antibodies, thereby markedly enhancing stability and reducing mRNA leakage in human serum. Additionally, LNP modified with OH-PEG exhibited reduced immunogenicity, which was crucial for repeated drug administrations. The present work elucidated the crucial role of OH-PEG in evading human pre-existing anti-PEG antibodies, and discovered that the current pre-clinical studies inadequately simulated the biological effects of clinical pre-existing anti-PEG antibodies on such formulations through interspecies study, which had a profound impact on clinical translation of PEGylated nanomedicines.

## Introduction

Polyethylene glycol (PEG) is a popular molecule in the pharmaceutical field, commonly used as an excipient to substantially increase the hydrophilicity of precursor.^1^ Conjugating PEG on the surface of nanocarriers or with proteins, which is called “PEGylation”, would reduce recognition by circulation opsonin and subsequent clearance by mononuclear phagocyte system (MPS), prolonging their circulating half-life *in vivo*.^2^ Meanwhile, PEGylation on nanocarriers is capable of enhancing colloidal stability by steric repulsion^3^ and managing passive targeting at tumor tissues through the enhanced permeability and retention (EPR) effects.^4^ Numerous PEGylated nanocarriers have been marketed or in clinical trials such as Doxil^®^, Onpattro^®^, and especially two universally inoculated LNP-mRNA vaccines for SARS-CoV-2, Pfizer BNT162b2 and Moderna mRNA-1273.^5^

Contrary to the traditional idea that PEG is non-immunogenic, an increasing number of studies reported detection of anti-PEG antibodies in animals and human.^6–8^ These antibodies rapidly recognize and bind to PEGylated nanocarriers, activate the complement system, leading to accelerated blood clearance of PEGylated nanocarriers.^9,10^ Additionally, the activated complement proteins would possibly mediate complement related hypersensitivity reactions and premature leakage of encapsulated drugs, significantly affecting the use of PEGylated nanomedicines.^11,12^ More worriedly, pre-existing anti-PEG antibodies have been detected in healthy individuals and clinical patients who have not been exposed to PEGylated therapeutics.^13^ The mechanism behind the formation of pre-existing anti-PEG antibodies is currently ambiguous but may be closely related to the widespread use of PEG as an additive in cosmetic products^14^ and PEG-based pharmaceutical excipients in intravenous preparations.^15^ Over time, the prevalence of pre-existing anti-PEG antibodies in the population has shown an increasing trend, rising from 0.2% in 1984^16^ to over 90% in 2019.^17^ In addition to its high prevalence, recent studies have discovered an alarming issue that the presence of pre-existing anti-PEG antibodies would hamper the efficacy and safety of PEGylated nanocarriers including activating the complement system,^18^ increasing blood clearance rate of PEGylated nanocarriers^19^ and causing injection reactions and other hypersensitivity reactions (HSRs). For example, Kozma *et al.* found association between extreme levels of pre-existing anti-PEG antibodies and an increasing risk for HSRs or anaphylaxis to LNP-mRNA vaccines.^20,21^ Khalil *et al.* discovered a high prevalence of pre-existing anti-PEG IgG (13.9%) and IgM (29.1%) in 701 children ALL patients receiving PEG-asparaginase and reduced asparaginase activities in an antibody-concentration dependent manner.^22^

Optimizing the reactogenicity of PEG with different terminal groups is one of the possible strategies to reduce the influence of clinical pre-existing anti-PEG antibodies.^23^ The PEG terminus in most marketed nanomedicines is methoxy groups (MeO-PEG), such as MeO-PEG-DSPE used in Doxil^®24^ and Onivyde^®^,^25^ MeO-PEG-DMG used in Onpattro^®26^ and Spikevax^®^,^23^ and ALC-0159 used in Comirnaty^®^.^27^ One reason for the universal use of MeO-PEG is that it has a simple structure with an inert methoxy group at one end, allowing for minimal crosslinking during synthesis and ensuring high homogeneity of the final products. Previous studies revealed that the structure of synthetic anti MeO-PEG antibodies would not be suitable for binding with PEG of hydroxy group (OH-PEG)^28^ and the immunogenicity of OH-PEG-DSPE modified liposomes was lower than MeO-PEG-DSPE in mice.^29^ However, they paid little attention to binding differences of human pre-existing anti-PEG antibodies to PEG with different termini and potential biological effects mediated by human pre-existing antibodies.

In the present study, pre-existing anti-PEG antibodies in 2074 human blood samples were analyzed on prevalence, sex and age correlation, and the binding affinities towards PEG with various termini were detected. Human pre-existing anti-PEG antibodies were found to be predominantly terminus-selective and OH-PEG exhibited the weakest binding among MeO-PEG, NH_2_-PEG, and COOH-PEG. Hence, OH-PEG modified LNP (OH-LNP) was synthesized and showed evasion of recognition by pre-existing anti-PEG antibodies compared with MeO-PEG modified LNP (MeO-LNP), therefore efficiently reduced complement activation. To explore whether OH-LNP could improve the biological function of LNP induced by pre-existing anti-PEG antibodies, drug leakage, cell uptake by human hepatocytes and macrophages were investigated, and OH-LNP demonstrated overall advantages over MeO-LNP. In addition, OH-PEG conjugates modified different nanocarriers like liposomes and micelles were also found to efficiently evade pre-existing anti-PEG antibody recognition. Considering the significance of pre-clinical studies, interspecies difference of selectivity of anti-PEG antibodies was examined on mice, rats and dogs. Surprisingly, it turned out that anti-PEG antibodies induced in animal models showed selectivity towards PEG repeated unit; meanwhile, OH-PEG modified nanocarriers could not evade complement activation in pre-stimulated animal models, indicating that preclinical studies were unable to simulate real clinical scenarios. To sum up, our work discovered the PEG terminus selectivity of human pre-existing anti-PEG antibodies and identified OH-PEG as solution to evade clinical pre-existing anti-PEG antibodies for efficient delivery of LNP.

## Results and discussion

### Methodology refinement of anti-PEG antibodies detection

The specificity of the secondary antibodies was first evaluated through a cross-reactivity test. As shown in Supplemental Figure 1, the secondary antibodies specifically bound to their corresponding targets. Tween-20 (polysorbate) is a widely applied non-ionic detergent in antibodies detection, which was also utilized in this work to reduce non-specific binding. However, some studies have reported that Tween might interfere with the detection of anti-PEG antibodies and suggested that using detergents without repeating ethylene glycol units, such as CHAPS, could result in higher OD values.^30^ In fact, there were literatures proving that Tween-20 did not interference with anti-PEG antibodies detection in pig blood, due to the entirely different epitope exposure between short-branched PEG in Tween and long-linear PEG in MeO-PEG-DSPE.^31^ To refine the choice of detergents in ELISA protocols, we firstly utilized different concentrations of Tween-20 to evaluate the impact of Tween as washing buffer on anti-PEG antibodies detection in human serum. As shown in Supplemental Figure 2, there were no significant differences between 0.05% and 0.1% Tween-20 when MeO-PEG-DSPE was employed as antigen to detect pre-existing anti-PEG antibodies. Furthermore, we incubated a series of concentrations of Tween-20/PBS (v/v) immediately after the blood sample or commercial anti-PEG antibodies incubation step to further confirm whether Tween would compete with the coating antigen for binding to the anti-PEG antibodies. As shown in Figure 1, incubation with no more than 0.05% Tween-20/PBS (v/v) would not affect the detection of anti-PEG antibodies which generated from mouse, rat and dog by pegylated liposome stimulation, and commercial anti-PEG antibodies. Although incubating with more than 0.05% Tween-20/PBS (v/v) would influence the detection of anti-PEG antibodies in animal serum and commercial antibodies, it would still not interfere with the binding of human pre-existing anti-PEG antibodies to MeO-PEG-DSPE. At the same time, We used human standard IgM and IgG antibodies (not anti-PEG type) were utilized to test whether the higher signals produced by using CHAPS as wash buffer were generated by anti-PEG antibodies. According to previous studies, the total content of immunoglobulins in human serum (7–16 mg/mL for IgG and 0.5–2 mg/mL for IgM) is over 10,000 times higher than the concentration of pre-existing anti-PEG antibodies (52 ng/mL for IgG and 22 ng/mL for IgM).^32^ Therefore, the ability of two detergents to eliminate non-specific binding was tested under conditions where the total IgM and IgG concentrations were 100 times and 1,000 times greater than those of the anti-PEG antibodies (see methods). As shown in Supplemental Figure 3, the addition of standard human antibodies had no effect on the detection of anti-PEG antibodies when using 0.05% Tween-20/PBS as the washing buffer, indicating its powerful ability to remove nonspecifically bound antibodies. However, when 0.05% CHAPS/PBS was used as the washing buffer, the addition of standard antibodies significantly increased the detection signal of anti-PEG IgG, suggesting that common IgG would also bind nonspecifically to the plate and be detected by the secondary antibodies using CHAPS/PBS as washing buffer. Even increasing the concentration of CHAPS to 0.1% could not completely eliminate non-specific binding by human standard IgG (Supplemental Figure 4). In addition to testing the ability of CHAPS to eliminate non-specific binding of the anti-PEG antibodies (which can be considered the primary antibody), its ability to eliminate non-specific binding of the secondary antibody was also evaluated by adding goat anti-human IgM or IgG (HRP) or PBS to blank plates. As shown in Supplemental Figure 5, due to the absence of corresponding antigen and primary antibodies on the plate, Tween-20 completely removed the non-specific binding of secondary antibody. However, the OD values in the CHAPS group were significantly higher than those in the PBS control, indicating that there was still a substantial amount of non-specific binding of HRP-labeled secondary antibodies in blank plates. At last, we decided 0.05% Tween/PBS instead of CHAPS/PBS was more capable of serving as detergents in ELISA detection of anti-PEG antibodies.

**Figure 1.**
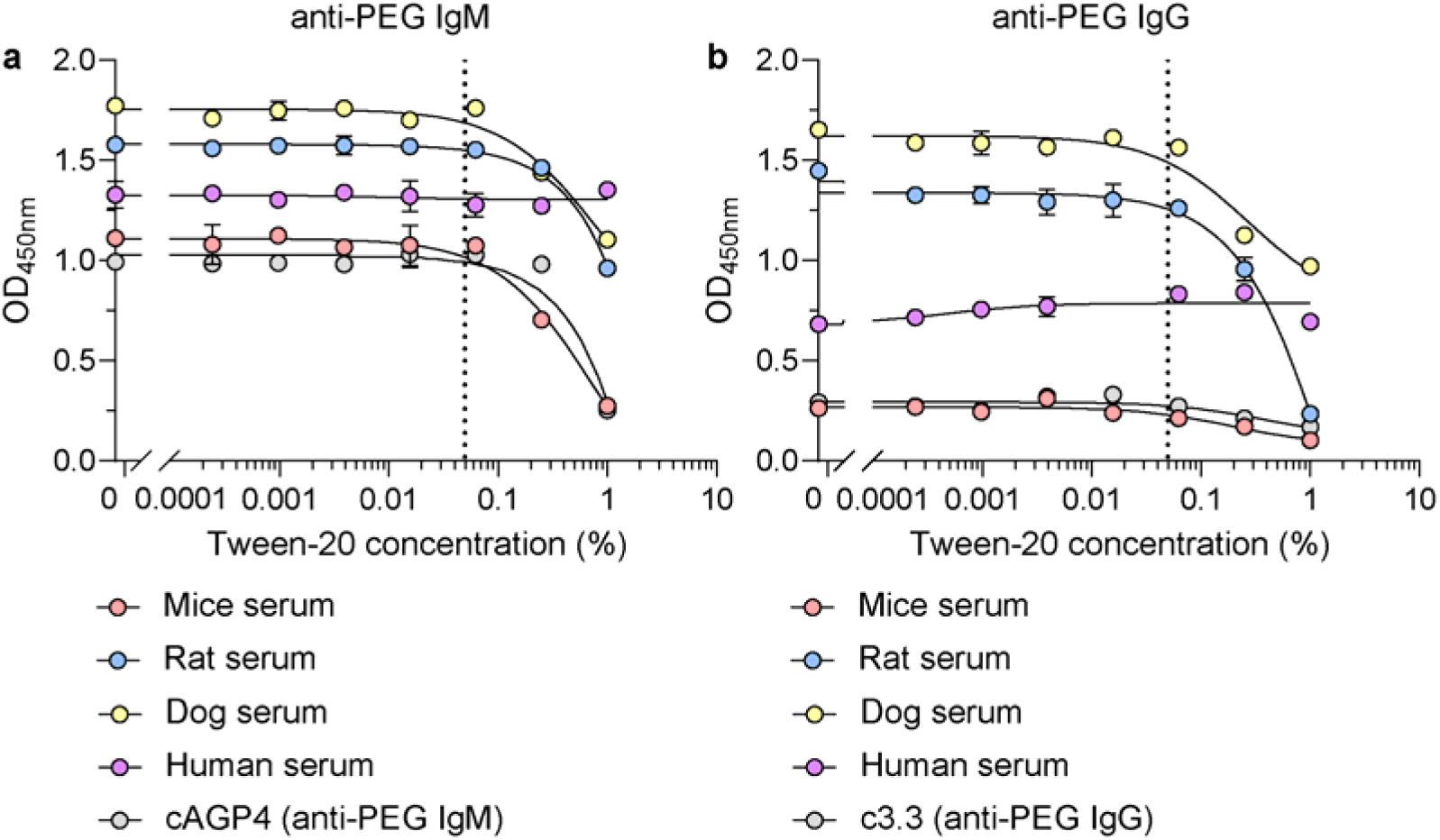
Effect of Tween-20 on anti-PEG antibody detection. After incubated with different concentration of Tween-20, levels of anti-PEG IgM (a) and IgG (b) were evaluated by ELISA assay. Data are means ± SDs (n = 2).

### Prevalence of human pre-existing anti-PEG antibodies

Human peripheral serum samples were collected from a pool of 2074 healthy donors from Jing’an District Central Hospital (Shanghai, China), aging from 16 to 95 with a female-male ratio 1.66. Direct sandwich ELISA (see Methods) was performed to measure the positive rates of pre-existing anti-PEG IgM and IgG using MeO-PEG-DSPE (the PEG molecular weight was 2000 Da) as the antigen and the preset lower limit value was 2.1 times of the control buffer (5% BSA) group for titration calculation (as previously described)^33^. Due to the uncertainty of structure, binding affinity, and pattern of the pre-existing anti-PEG antibodies in human samples, we arbitrarily deemed OD_450nm_>1.0 (1:4 dilution of the human serum sample, the same below) as strongly positive, 1.0>OD_450nm_>0.5 as positive (gray zone) and OD_450nm_<0.5 as negative (Figure 2a) to briefly distinguish the antibody level. Similar to previous reports,^33^ we found anti-PEG IgM was more prevalent in population, with a ratio of 18.0% for the strongly positive and 27.5% for the positive. While anti-PEG IgG only accounted for 0.8% for the strongly positive and 18.1% for the positive. The specificity of the ELISA methods was verified by inhibition study using PEGylated liposome (see the preparation in Methods) as competitive binding agent. As shown in Supplemental Figure 6, the addition of liposomes led to 90% reduction of the absorbance values, suggesting the vast majority of the detected signal originated from the pre-existing anti-PEG antibodies. To verify the reasonability of our definition, full dilutions of anti-PEG IgM of 120 samples from three gradations were prepared (Figure 2b) and the correlation between their OD_450nm_ (1:4 dilution) values and IgM titers was analyzed (Figure 2d). Anti-PEG antibody titer was estimated by end-point dilution in the ELISA by analyzing duplicate serial 4-fold dilutions from 1:4 to 1:65536. As shown in Figure 2c, IgM titer of the strongly positive group was remarkably higher than that of the positive and negative groups (p<0.001) and IgM titer of the positive group was also significantly higher than that of the negative group as expected (p<0.001), exhibiting prominent variation among groups. Also, the OD_450nm_ values of 1:4 diluted samples were of good linear correlation with their IgM titers (Figure 2d), validating the former as a suitable indicator for large population screening. The antibody titers in negative and strongly positive human samples which we arbitrarily deemed were evaluated via the fitted curve, and found that the IgM titers in negative samples with OD<0.5 and strongly positive samples with OD>1 were below 1:50 and above 1:100. However, the linear correlation between OD_450nm_ values and titers of anti-PEG IgG was not established (Supplemental Figure 7), possibly due to their relatively low OD_450nm_ values leading to inaccurate titers. In addition, we analyzed the prevalence of anti-PEG antibodies in human by sex and age (Supplemental Figure 8). Consistent with other studies,^34^ anti-PEG antibodies were more prevalent in females (IgM 21.8% *vs.* 11.7%; IgG 1.0% *vs.* 0.4%, p<0.0001) and the antibody level decreased with age.

**Figure 2.**
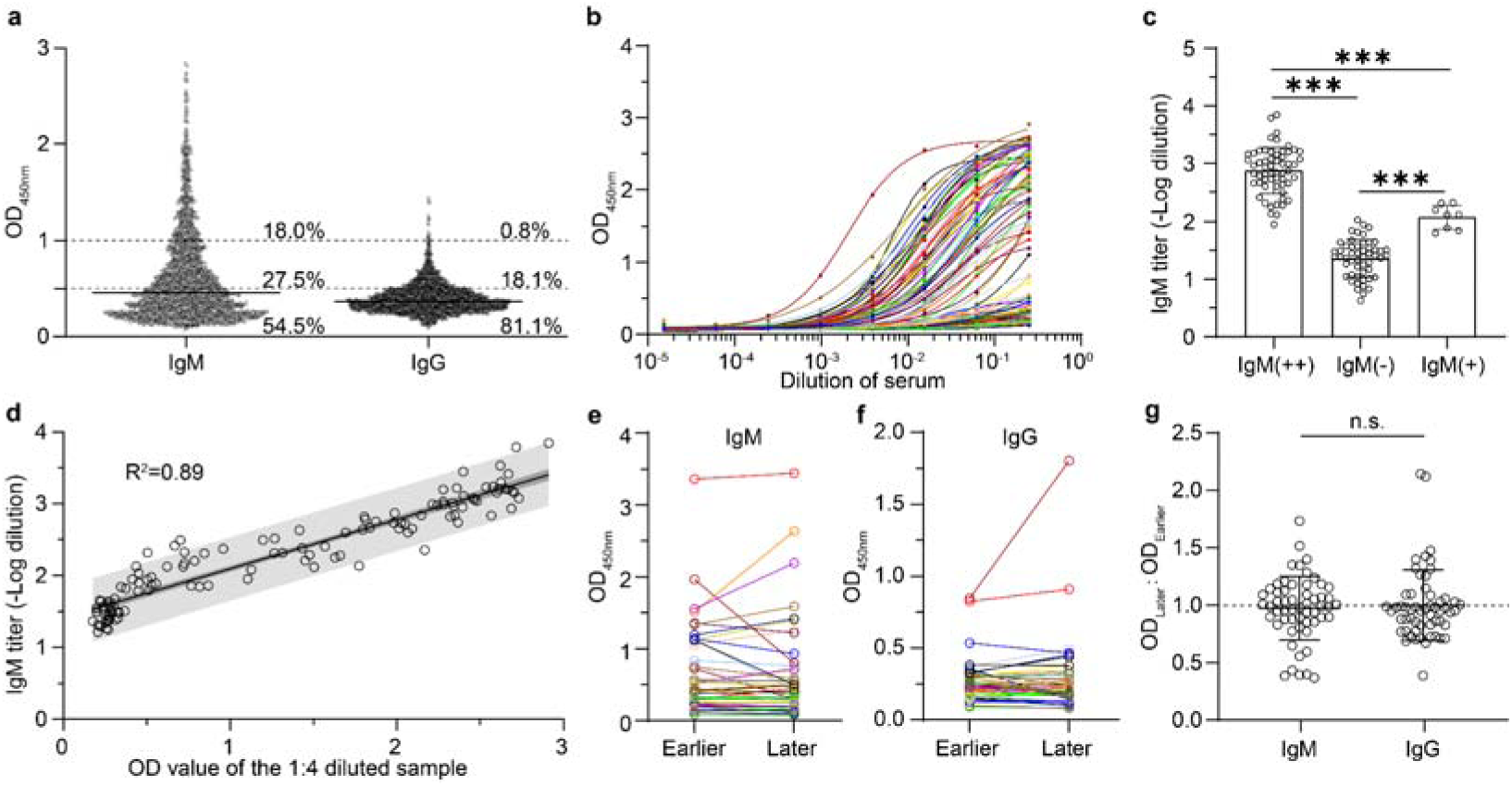
Prevalence of human pre-existing anti-PEG antibodies. **(a)** Levels of anti-PEG IgM and IgG in all donors (n=2074), using MeO-PEG-DSPE as antigen and OD_450nm_ values of 4-times dilution of human serum as measurement standard. Full dilutions **(b)** and titrations **(c)** of pre-existing anti-PEG IgM in 125 human samples. Titration was calculated as -log(dilution, whose OD_450nm_ was 2.1 times of the control buffer) using the embedded equation of specific binding in GraphPad Prism 9.0. **(d)** The relationship between single point OD_450nm_ values and titrations of anti-PEG IgM. Linear regression line (Y = 0.6841*X+1.412, R^2^=0.89) was shown in solid line. Grey area showed 95% prediction intervals. Levels of anti-PEG IgM **(e)** and IgG **(f)** in samples of two time-points (at least 3 months interval) from a single donor. **(g)** The variation of anti-PEG IgM and IgG, calculated by OD_Later_:OD_Earlier_ (n=55). Data are means±SDs. Statistical significance is evaluated using Student’s t-test with GraphPad Prism 9.0. (n.s. indicates non-significant, ∗∗∗*p<0.001*).

Furthermore, 55 human serum samples (two time-points from a single donor with a ≥3 months interval) were selected to confirm if the pre-existing anti-PEG antibodies were transient or long-lasting. As shown in Figure 2e and Figure 2f, both pre-existing anti-PEG IgM and IgG were minimally fluctuant. Also, the comparison between variations of anti-PEG IgM and IgG (Figure 2g) showed no significant difference (utilizing OD_Later_:OD_Earlier_ as the index). The fact that pre-existing anti-PEG antibodies were permanent indirectly indicated the potentially long-lasting impacts of these antibodies on PEGylated nanocarriers.

### PEG terminus selectivity of human pre-existing anti-PEG antibodies

It was reported that binding affinities of anti-PEG antibodies with PEG of distinct terminal groups were quite different.^23^ We selected 40 serum samples containing varied levels of anti-PEG IgM (Figure 3a) and IgG (Figure 3d) to detect their binding with PEG-DSPE (with a PEG molecular weight of 2000 Da) with different termini including methoxy (MeO), amino (NH_2_), carboxy (COOH) and hydroxy (OH) utilizing the same ELISA protocols but different antigens (see Methods). The heat map clearly indicated that both human anti-PEG IgM and IgG bound with PEG-DSPE in a PEG termini-dependent manner. Among those, the hydroxy terminus of PEG (OH-PEG-DSPE) apparently exhibited the least reactogenicity to human pre-existing anti-PEG IgM and IgG. Furthermore, we expanded the measurement to all human serum samples which were strongly positive for anti-PEG IgM (n=373, Figure 3b) and IgG (n=16, Figure 3e) observed in this work to compare the whole positive rate of human serum samples against MeO-PEG-DSPE and OH-PEG-DSPE. The result showed that only 6% anti-MeO-PEG IgM and 6% IgG strongly positive samples were strongly positive when using OH-PEG-DSPE as antigen. Also, 155 IgM and 8 IgG positive samples from Fudan University affiliated Shanghai Pudong Hospital^33^ (Shanghai, China) were also detected for PEG terminus selectivity (Supplemental Figure 9). The result showed similar trend with 10% anti-MeO-PEG IgM and 0% IgG strongly positive samples were strongly positive when using OH-PEG-DSPE as antigen. The parallel results of samples from two different centers further verified the evasion effect of OH-PEG-DSPE to pre-existing anti-PEG antibodies. Considering the stringency of the detection, we also tested over 300 human serum samples which were anti-PEG antibodies negative for binding with OH-PEG-DSPE (Figure 3c and 3f). The result showed almost all anti-MeO-PEG IgM and IgG negative samples were anti-OH-PEG antibodies negative. Despite that PEG-DSPE was commonly used for construction of PEGylated liposomes, PEG materials with different hydrophobic groups were appended into synthesis of other common PEGylated nanomedicines. PEG-DMG (for preparation of LNP), PEG-PDLLA (for preparation of polymeric micelles) and PEG-PLGA (for preparation of PLGA-nanoparticles) were employed to confirm whether other OH-PEG materials could evade pre-existing anti-PEG antibodies. As shown in Supplemental Figure 10, OH-PEG with different PEG chain lengths or conjugated hydrophobic groups maintained weaker binding with pre-existing anti-PEG IgM.

**Figure 3.**
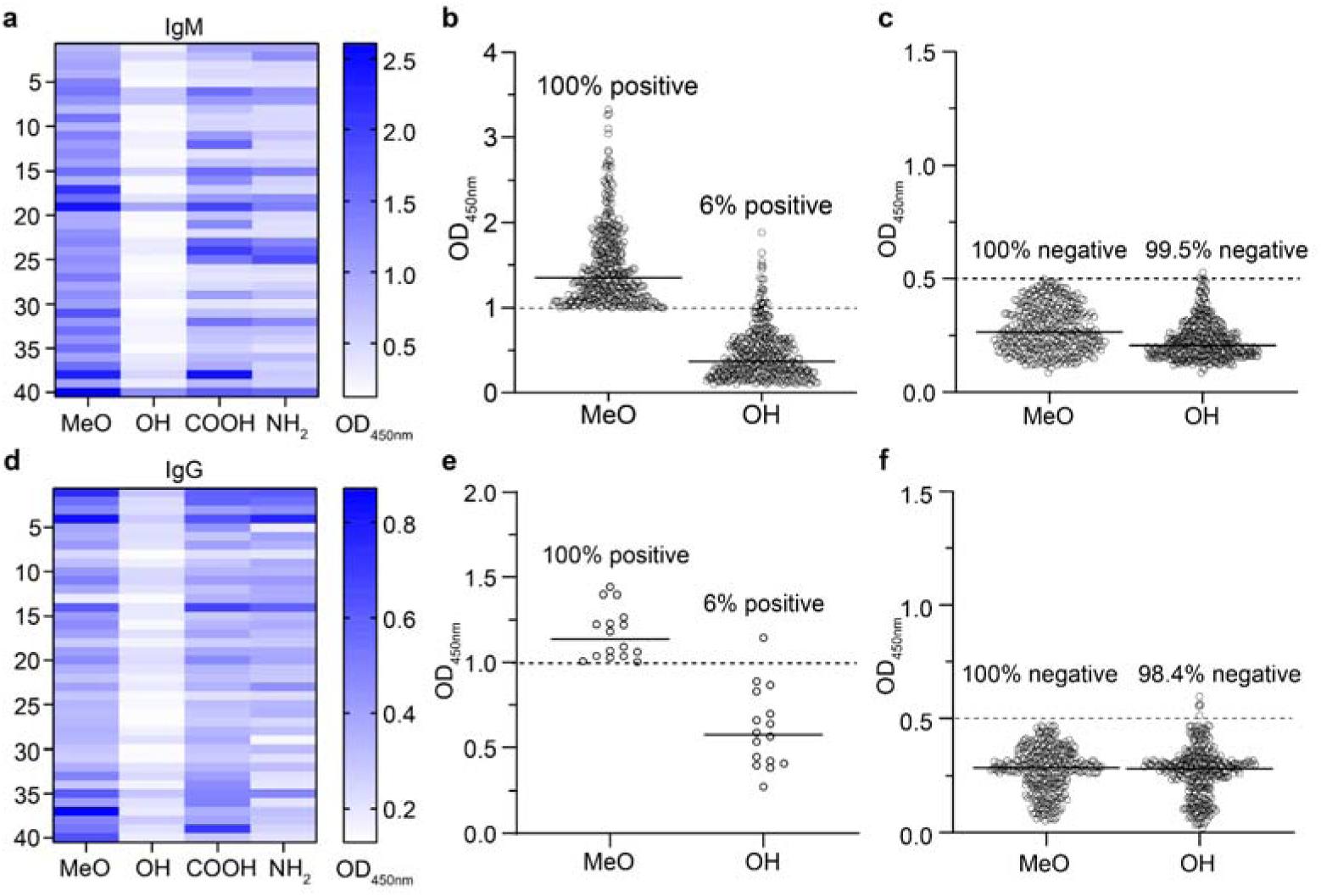
PEG terminus selectivity of human pre-existing anti-PEG antibodies. The binding of anti-PEG IgM **(a)** and IgG **(d)** to PEG-DSPE with different termini (the ordinates represented each human serum sample tested and the abscissas represented PEG-DSPE of different termini examined) (n=40). The positive rate of anti-PEG IgM **(b)** or IgG **(e)** in strongly positive human samples against OH-PEG-DSPE (n=373 and 16). The positive rate of anti-PEG IgM **(c)** or IgG **(f)** in negative human samples against OH-PEG-DSPE (n=350 and 308). Data are shown in dots and means, and analyzed with GraphPad Prism 9.0.

To further confirm the ability of OH-PEG to evade pre-existing anti-PEG antibodies, isothermal titration calorimetry (ITC) was utilized to directly quantify the interaction between different PEG materials and anti-PEG antibodies in serum^35^. The measurements were taken under constant temperature conditions by titrating PEG-DSPE into serum (see methods). MeO-PEG was titrated to anti-PEG antibodies positive (stimulated by intravenously injecting 2.5 mg HSPC/kg PEGylated liposome in rats 6 days ago) or negative (wild-type) rat serum to determine if ITC was capable of detecting binding of anti-PEG antibodies to PEG materials. As shown in Supplemental Figure 11, the heat release from titration of anti-PEG antibody positive rat serum with MeO-PEG was significantly higher than that from the titration of negative serum. As the antibody in sample cell became saturated with PEG materials, the heat signal gradually decreased, whereas the heat release in antibody-negative rat serum was minimal and did not vary with the increase of PEG materials concentration. These results indicated that ITC could distinguish between antibody-positive and antibody-negative serum.

ITC was further used to measure the heat release from titration of MeO-PEG and OH-PEG into human serum. As shown in Supplemental Figure 12, human serum samples identified as antibody negative by ELISA (OD<0.5) showed weak reactions with both PEG materials, with low heat release and no trend of decreasing heat release as the proportion of PEG in the sample cell increased. In contrast, titration of MeO-PEG in all antibody positive sera (OD>1) released significantly more heat than OH-PEG, and the heat release gradually decreased as the reaction approached saturation. The estimated dissociation constant (K_D_) ranged from 3 to 15 µM. While no significant heat release or specific binding were observed during titration of OH-PEG in antibody positive serum, thus the KD values could not be fitted (Supplemental Table 1). These results directly validate the interaction between human pre-existing anti-PEG antibody with MeO-PEG, but not OH-PEG. It also validated the effectiveness of our ELISA method for detecting the presence of pre-existing antibodies and the binding between different PEG materials and pre-existing anti-PEG antibodies. To the best of our knowledge, this was the initial report on PEG terminus selectivity of human pre-existing anti-PEG antibodies. We discovered that the binding affinities of OH-PEG with human pre-existing anti-PEG antibodies were significantly lower than MeO-PEG, which was the only approved PEG terminus in the preparation of PEGylated nanomedicines. It was anticipated that the application of OH-PEG conjugates in nanocarriers was a feasible approach to evade recognition and side effects of human pre-existing anti-PEG antibodies.

### Hydroxy PEG efficiently evades pre-existing anti-PEG antibodies recognition of LNP

To further investigate the effect of PEG terminus on lipid nanoparticles (LNP), PEG-DMG (1.5%, 3%, or 5% molar ratio to the total lipids) with hydroxy or methoxy terminus was used for the preparation of LNP by the microfluidic mixing method (see Methods) and characterized by dynamic light scattering (DLS). As shown in Supplemental Table 2, the particle size of LNP gradually decreased as the proportion of PEG modification increased but remained within the range of 65-75 nm. ELISA was employed to assess the binding affinities between LNP and pre-existing anti-PEG antibodies in human serum, using LNP modified with DMG-PEG as antigens (see Methods). As shown in Figure 4a-c, LNP modified with OH-PEG-DMG (OH-LNP) showed lower binding with pre-existing antibodies across various modification ratios compared to LNP modified with MeO-PEG-DMG (MeO-LNP). Due to the hydrophilicity of PEG, the increase in PEG modification ratio weakened the non-specific binding, resulting in more significant difference in specific binding between methoxy and hydroxy PEG modification. After entry blood stream, plasma proteins rapidly deposited on the surface of nanoparticles and formed protein corona, which significantly influenced the *in vivo* fate of nanomedicines.^36^ Immunoglobulin was one of the most important opsonin in protein corona. Therefore, the protein corona on the surface of LNP was separated and the adsorbed plasma proteins were characterized by SDS-PAGE and western blot assay to evaluate the interaction of pre-existing anti-PEG antibodies with LNP. After incubation with the anti-PEG antibodies negative human serum samples, neither OH-LNP nor MeO-LNP showed significant deposition of IgM. While in anti-PEG antibodies positive human serum samples, OH-LNP demonstrated significant lower antibody binding levels (Figure 4d-f). Even though the overall concentration of IgG in human serum was higher than that of IgM (Figure 4d), LNP exhibited greater adsorption of IgM, suggesting that PEG-specific antibodies were predominant of the IgM isotype, consistent with the lower detection rate of anti-PEG IgG and the higher level of anti-PEG IgM observed in Figure 2a. We further detected the binding of pre-existing anti-PEG antibodies with other OH-PEG modified nanocarriers, including liposomes (sLip), PLGA nanoparticles (NP) and polymer micelle (PM) (see the preparation in Methods). As expected, the three types of nanocarrier modified with OH-PEG exhibited weaker binding with PEG antibody compared to those with MeO-PEG (Supplemental Figure 13).

**Figure 4.**
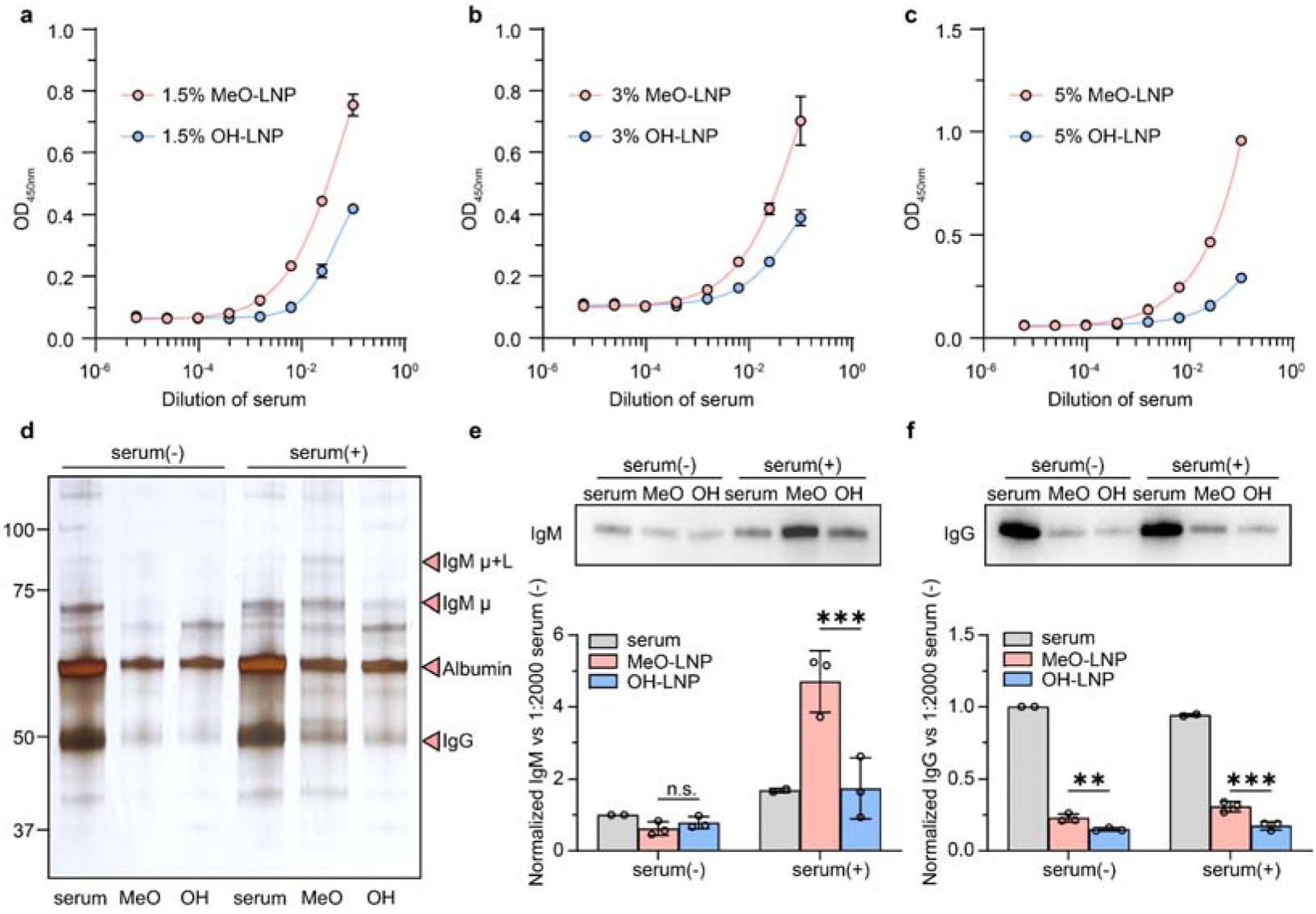
Hydroxy PEG efficiently evades the recognition of LNP by human pre-existing anti-PEG antibodies. The binding of pre-existing anti-PEG IgM to LNP modified with 1.5% **(a)**, 3% **(b)**, or 5% **(c)** OH-PEG-DMG or MeO-PEG-DMG. **(d)** Separation of the protein corona by SDS-PAGE and staining with silver. Western blot detection and quantification of anti-PEG IgM **(e)** and IgG **(f)** on the surface of LNP after incubation with antibody negative serum (serum(-)) or positive serum (serum(+)) for 1 h. Data are means±SDs (n=3). Statistical significance is evaluated using Two-way ANOVA with GraphPad Prism 9.0 (n.s. indicates non-significant, ∗∗*p*<0.01, ∗∗∗*p<0.001*).

### Functional effects of human pre-existing antibodies on LNP

To explore whether the evasion capability of OH-LNP towards PEG pre-existing antibodies could improve the biological functions, complement activation was detected in human serum by western blot. The dissociation of complement protein 3 (iC3b:C3 ratio) was used as the symbol of complement activation, because C3 was the most abundant complement proteins in plasma and played a significant role in subsequent complement activation as the core of three complement activation pathways.^37^ As shown in Figure 5a and 5b, LNP did not induce significant complement activation in anti-PEG antibody negative human serum. However, in anti-PEG antibody positive human serum, MeO-LNP induced significant C3 cleavage compared with OH-LNP, which indirectly supporting the validity of using OD values to distinguish between antibodies negative and positive serum. Human serum C5a level was also measured using an ELISA kit (ab193695) after incubation with LNPs, following the manufacturer’s instructions. As shown in Supplemental Figure 14, the trend paralleled C3 cleavage, indicating that MeO-LNP induced significant complement activation in antibody-positive human serum compared to OH-LNP. The endpoint of complement activation was the formation of the membrane attack complex (MAC) on the surface of nanoparticles, disrupting the integrity of the lipid membrane and resulting in the premature release of the encapsulated cargo.^38^ Due to the interference of RNA fragments in serum with RiboGreen™ dye, the impact of PEG antibodies and complement activation on mRNA leakage was investigated using RT-qPCR. After incubation with PEG antibody positive human serum at 37 _ for 2 h in the presence of RNase, the total eGFP-mRNA payload of OH-LNP remained unchanged; while in MeO-LNP, almost half of the encapsulated mRNA disappeared due to the enhanced complement activation capability (Figure 5c). The cellular uptake of LNP was assessed in human hepatocytes LO2 and THP-1 derived macrophages cell lines after incubation with sera, aiming to characterize the impact of pre-existing anti-PEG antibodies on both targeting and off-targeting capabilities *in vitro*. As shown in Figure 5d and Figure 5e, the presence of anti-PEG antibodies did not influence the uptake of LNP by hepatocytes, but significantly increased the uptake of MeO-LNP by macrophages, which had a much higher antibody and complement binding capacity. To further confirm the important role of complement activation in LNP leakage and macrophage uptake, antibody positive human serum was pre-incubated at 56℃ for 30 min to dissociate heat-labile complement protein, or mixed with EDTA-PBS (10 mM) to deactivate complement system. The mRNA leakage and macrophage uptake were evaluated by RT-qPCR. As shown in Supplemental Figure 15a, the complement deactivation by both strategies significantly mitigated the damage of MAC on the integrity of the lipid membrane, resulting in reduced mRNA leakage. In contrary, as shown in Supplemental Figure 15b, the complement deactivation by heating nearly completely prevented macrophage uptake of MeO-LNP, while EDTA only partially reduced the macrophage uptake. One possible reason is that heating-induced complement dissociation is irreversible, whereas EDTA inhibits complement activation by chelating metal ions in the serum. However, once the serum/LNP mixture was introduced into the RMPI 1640 culture medium and incubated with cells, the presence of Ca^2+^ and Mg^2+^ in the medium might further facilitate complement activation by MeO-LNP. Regardless, OH-LNP demonstrated significantly superior effects in antibody positive human serum. The mechanisms behind anaphylaxis and other manifestations of hypersensitivity triggered by mRNA-LNP based SARS-CoV-2 vaccines or other PEGylated nanomedicines such as liposomal doxorubicin were considered to associate with complement activation. Moreover, C3 protein cleavage and an increase in C5a level were observed in untreated (Supplemental Figure 16, 17) or sterile PBS-treated anti-PEG antibodies positive human serum, indicating a potential hypersensitivity of the complement system, thus the use of MeO-PEG mediated nanomedicines in this population might pose excessive risks.

**Figure 5.**
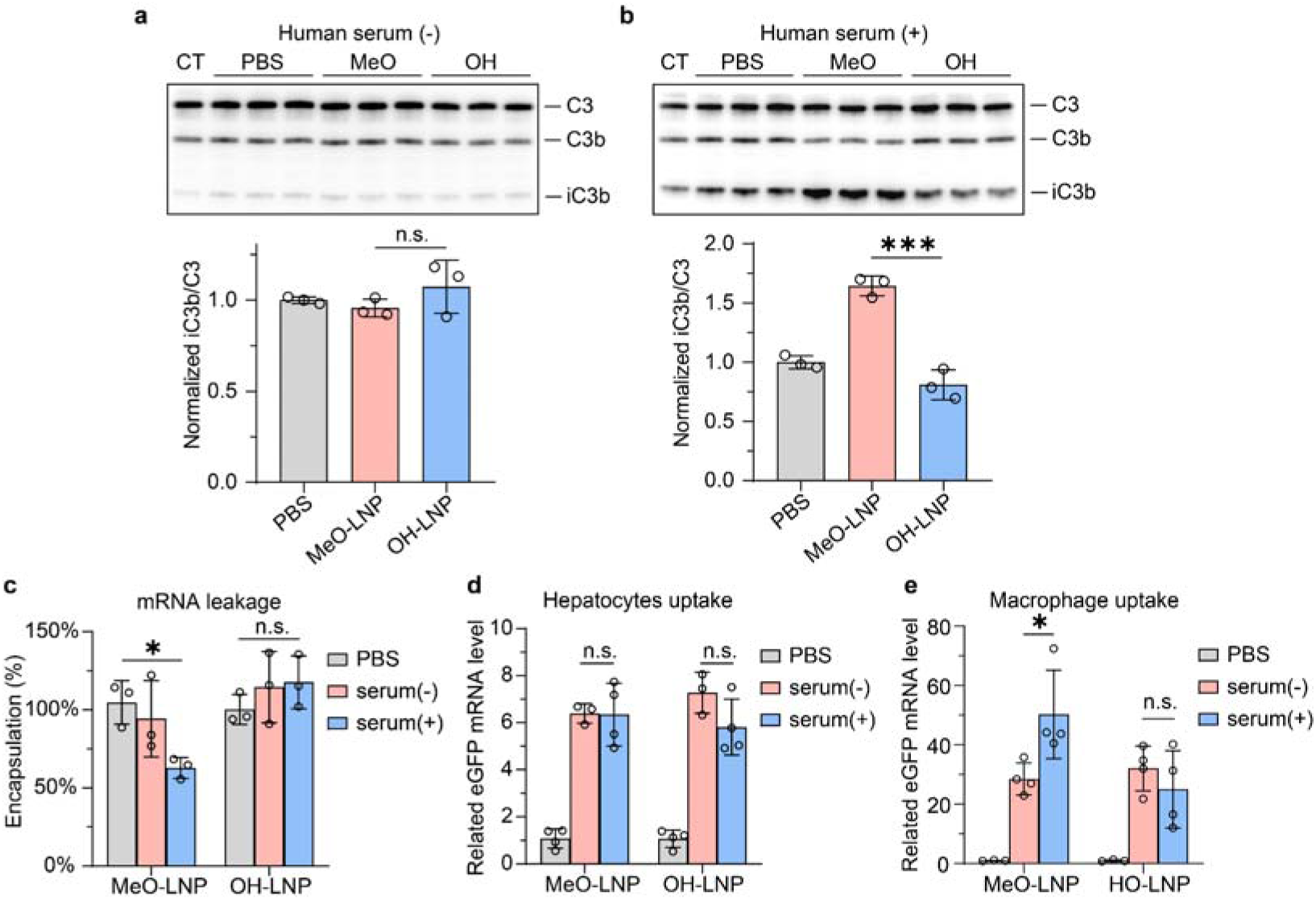
Biological functions of human pre-existing antibodies on LNP. Western blot detection and quantification of C3 cleavage in pre-existing anti-PEG antibody negative **(a)** or positive **(b)** human serum after incubation with LNP for 1 h. **(c)** The remaining eGFP mRNA encapsulated in LNP after incubation with antibody positive human serum for 2 h in the presence of RNase. Uptake of eGFP-LNP by human hepatocyte LO2 cells **(d)** and macrophage THP-1 cells **(e)** in the presence of anti-PEG antibody positive human serum, quantified by RT-qPCR. Data are means±SDs (n=3-4). Statistical significance is evaluated using Two-way ANOVA with GraphPad Prism 9.0 (n.s. indicates non-significant, ∗*p<0.05*, ∗∗∗*p<0.001*).

### Interspecies contradiction of anti-PEG antibodies

In addition to the human pre-existing anti-PEG antibodies, anti-PEG antibodies (mostly IgM) could also be produced by marginal zone B cells through a T cell-independent pathway after repeated injections of PEGylated nanocarriers in animals.^30^ Nowadays, limited by few knowledges on the production of clinical pre-existing anti-PEG antibody, a large number of preclinical animal experiments utilized anti-PEG antibodies stimulated by empty PEGylated nanoparticles injection (mostly liposome) to simulate pre-existing antibodies.^39–42^ To explore the comparability between anti-PEG antibodies induced by PEGylated nanocarriers in animals and human pre-existing antibodies and whether the evading capability of OH-PEG could also be effective in animal models, BALB/c mice (5 mg HSPC/kg), SD rats (2.5 mg HSPC/kg) and beagle dogs (1 mg HSPC/kg) were intravenously injected with empty PEGylated liposomes to generate anti-PEG antibodies. After 6 days, blood was sampled and binding between the stimulated anti-PEG antibodies and MeO-PEG-DSPE or OH-PEG-DSPE was examined (see Methods). Unlike human pre-existing anti-PEG antibodies, OH-PEG failed to evade the binding of the stimulated anti-PEG antibody in mice or rats, and only exhibited a mild evading effect in dogs (Figure 6a-d). The contradiction might arise from the potential differences in the binding mechanisms between anti-PEG antibodies stimulated by PEGylated nanocarriers in animals and the human pre-existing anti-PEG antibodies. As shown in Supplemental Figure 18, the anti-PEG antibodies produced in rats likely targeted the repeated ethylene glycol units within the PEG chain, showing no selectivity for PEG terminus. In contrast, the pre-existing anti-PEG antibodies in humans primarily recognized PEG terminus. The distinctiveness in the binding was supposed to partially derive from their origins. As stated before, human pre-existing antibodies were deemed to be caused by constant application of cosmetics (mostly), food and industry containing PEG, which was mostly short and methoxylated.^43^ However, MeO-PEG-DSPE used in liposomes contains more than 40 repeated units and is densely modified on nanocarriers.^44^ Further research is needed to understand why human pre-existing anti-PEG antibodies exhibit PEG-terminus selectivity. Meanwhile, OH-LNP even increased complement activation level in both anti-PEG antibody negative and positive mice and rats serum samples, and comparable complement activation level in dogs although there was a slightly weaker antibody binding affinity (Figure 6e-g), which might be attributed to the ability of hydroxy to directly split C3.^45^ Additionally, the adjacent hydroxy PEG on the surface of LNP might serve as 3- and 4-hydroxy groups of mannoses, which could bind with calcium-dependent carbohydrate-recognition domain in mannose-binding lectin, and induce complement activation via lectin pathway.^46,47^ It was interesting that OH-LNP did not significantly induce complement activation in antibody negative human serum (Figure 5a), but not in negative animal serum, possibly due to the significant differences between human and animal serum in the pathways of complement activation of nanoparticles. In animals, the lectin pathway and classical pathway were found to be the predominant pathways, whereas in humans without specific antibodies, the alternative pathway was the primary activation route, with minimal involvement of the lectin pathway.^48,49^ Considering the obvious inter-species dissimilarity in selectivity of anti-PEG antibodies and complement activation pathway, along with the necessity of animal experiments before clinical trials for most nanomedicines, the similar binding of anti-PEG antibodies to MeO-PEG and OH-PEG, coupled with the robust complement activation of OH-PEG in animals, might shift researchers’ focus towards MeO-PEG, considered inert, and hinder the application of OH-PEG in human studies and the further development of drugs for clinical use.

**Figure 6.**
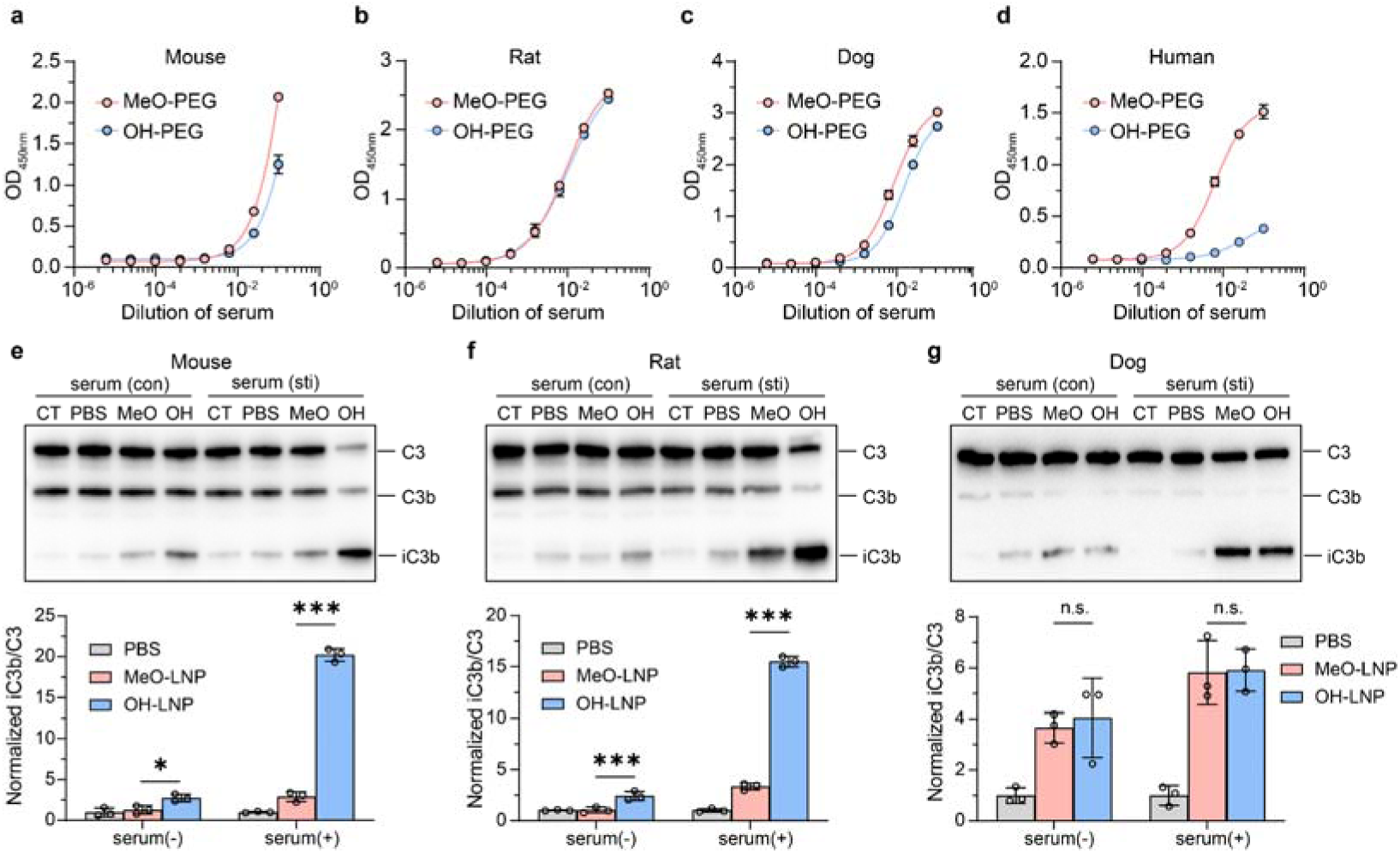
The Interspecies differences of evasion effect of OH-PEG on anti-PEG antibodies. The binding of OH-PEG and MeO-PEG with anti-PEG IgM pre-stimulated by sLip in mouse **(a)**, rat **(b)** and dog **(c)**, or pre-existing anti-PEG IgM in human serum **(d)**. Western blot detection and quantification of C3 cleavage in mouse **(e)**, rat **(f)** and dog **(g)** serum after incubation with LNP for 1 h (CT indicated ctrl serum without any incubation). Data are means±SDs (n=3). Statistical significance is evaluated using Two-way ANOVA with GraphPad Prism 9.0 (n.s. indicates non-significant, ∗*p<0.05*, ∗∗∗*p<0.001*).

### OH-LNP Immunogenicity among species

Considering that repeated injections of PEGylated nanocarrier could also lead to the production of corresponding anti-PEG antibodies, accelerate the clearance of subsequent doses and impact the clinical application of nanomedicines,^50^ the immunogenicity of OH-LNP was also investigated in different species. OH-LNP and MeO-LNP were intravenously injected into BALB/c mice (40 mg/kg lipid) or Beagle dogs (60 mg/kg lipid) weekly. One week after each injection, blood was sampled and anti-PEG antibodies were detected using MeO-PEG-DMG or OH-PEG-DMG as the antigen, respectively. As shown in Figure 7, repeated injections of MeO-LNP induced significant production of both anti-PEG IgM and IgG antibodies in beagle dogs, while OH-LNP group showed lower antibody levels at the first week and minimal detection of anti-PEG antibodies at the second week and third week. However, as shown in Supplemental Figure 19, neither type of LNP induced significant anti-PEG antibody production in mice, which might be related to the detachment of PEG-DMG from LNP for lack of enough antigen stimuli,^51^ and the much lower sensitivity and spectrum of symptoms in mice after receiving PEG nanomaterials compared to dogs, in which even a very small dose of PEGylated nanomaterials elicits strong immune reactions.^52,53^ These results suggested that OH-PEG modification not only evaded the binding of human pre-existing anti-PEG antibodies, but also reduced immunogenicity compared to MeO-PEG, manifesting its overall advantages in the construction of nanomedicines.

**Figure 7.**
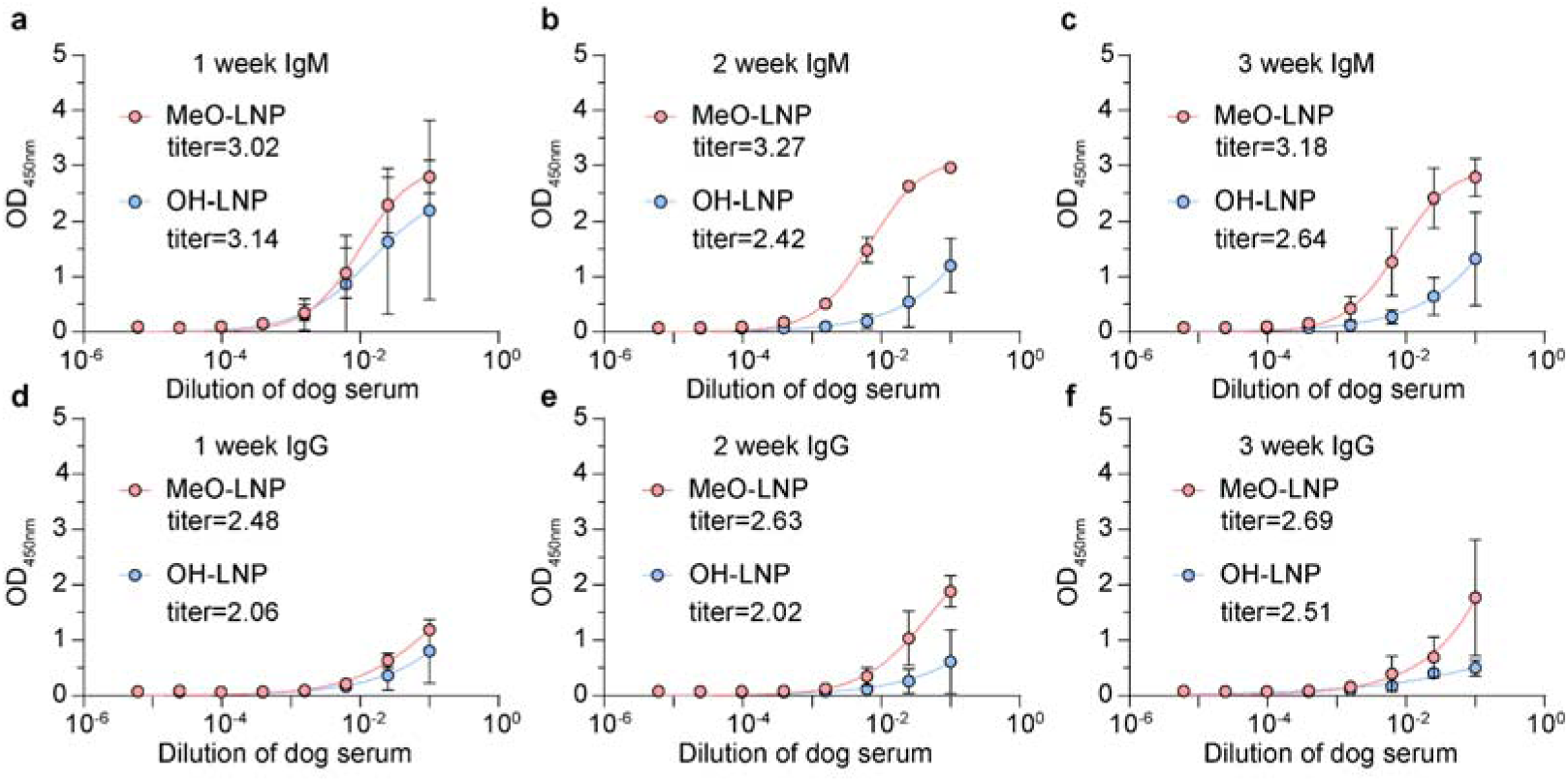
The immunogenicity of LNP in dogs. **(a-f)** The anti-PEG IgM and IgG was detected in dog serum from 1 week to 3 weeks after intravenous injection of MeO-LNP or OH-LNP weekly. Titer is -log (end-point dilution). Data are means±SDs (n=3) and analyzed with GraphPad Prism 9.0.

## Conclusion and discussion

In this study, we investigated the presence of pre-existing anti-PEG antibodies in a cohort of 2074 healthy donors, and delved into their prevalence. It was concluded that 18.0% and 0.8% of total population possessed high level of anti-PEG IgM and anti-PEG IgG antibodies, respectively. The level of pre-existing antibodies was related with gender and age. The relative higher rates of anti-PEG antibodies positive samples in female and middle-aged adults were probably due to more frequent usage of cosmetics with PEG as emulsifier and thickener. The persistence of pre-existing anti-PEG IgM and IgG antibodies was confirmed through a longitudinal observation of 55 samples over a three-month period. This sustained presence might be attributed to the regular use of cosmetics and toiletries containing PEG derivatives.

The most widely used approach to estimate unknown antibody levels is titration, which involves measuring the dilution at which the antibody binding signal reaches the baseline or a predetermined lower limit value. Our results in Figure 2d showed good linear correlation between antibody titers and OD values (R^2^=0.9), which proved that OD values were appropriate to represent the antibody levels. Due to the absence of chimeric anti-PEG antibodies with exactly the same structure, binding affinities, and patterns as pre-existing anti-PEG antibodies in human samples, in our opinion, no chimeric antibodies were suitable to serve as standard antibodies. Steve R. Roffler *et al*.^34,54^ used chimeric antibodies with mouse variable regions and human heavy chains as standard antibodies to quantify pre-existing anti-PEG antibodies, and found a significantly higher concentration of anti-PEG IgG compared to IgM, which contrasted sharply with the findings from our research and other research groups^21,55–58^. This paradox may stem from the inappropriate selection of measurement assays. They utilized chimeric antibodies (AGP4 for IgM and c3.3 for IgG) with Fab fragments generated in mice, without assessing the actual binding affinity of clinically pre-existing antibodies. Notably, the binding affinity of AGP4 was considerably weaker compared to c3.3. Therefore, if pre-existing anti PEG IgG and IgM in human serum sample have similar binding affinities to PEG, the IgG standard curve under identical measurement conditions will exhibit higher response values, resulting in higher detected IgG concentrations compared to IgM.

Through the measurement of the binding of pre-existing anti-PEG antibodies with various PEG materials via ELISA and ITC, our findings revealed that human pre-existing anti-PEG antibodies predominantly exhibited selectivity for PEG termini. OH-PEG exhibited the lowest binding with human anti-PEG antibodies among other commonly used PEG structures. The use of OH-PEG reduced the positive ratio by more than 90% and did not introduce any additional positive samples among those tested negatives for MeO-PEG, suggesting that introducing OH-PEG into nanocarriers is a viable strategy to evade pre-existing antibody recognition.

We investigated the bio-function of pre-existing MeO-PEG-specific antibodies on lipid nanoparticles and observed that the adsorption of anti-PEG antibodies on MeO-LNP triggered significant complement activation, resulting in mRNA leakage and unwanted macrophage uptake. In contrast, OH-LNP effectively reduced complement activation and preserved structural integrity by avoiding binding with human anti-PEG antibodies. It was reported that the delivery of LNP to liver was mostly mediated by albumin receptor and lipoprotein receptors,^59,60^ and there were no significant differences in the adsorption of albumin or lipoproteins such as APOE between MeO-LNP and OH-LNP observed on SDS-PAGE, indicating that the associated proteins might bind more to the lipid surface of LNP rather than the PEG chains.^61^ That might explain why anti-PEG antibodies did not affect the uptake of LNP by hepatocytes. To elucidate the influence of pre-existing anti-PEG antibodies on LNP, additional studies are needed to further investigate the dynamics of PEG-shedding and structural rearrangement, as well as lysosomal escape and gene expression in both targeting and off-targeting cells.

We found that in contrast to pre-existing anti-PEG antibodies in humans, anti-PEG antibodies induced by the injection of PEGylated nanocarriers in animals lacked selectivity for PEG terminal groups. Instead, they exhibited binding with repeated ethylene glycol units within the PEG chain. The differential binding patterns of PEG antibodies may depend on the specific antigenic stimuli. The generation of pre-existing anti-PEG antibodies in healthy individual was believed to result from the use of cosmetics containing large scale of PEG derivatives, in which the PEG derivatives with the highest concentration of use (such as laureth 4, C12-13 pareth 3, ceteth 9 and ceteareth 10) or with the greatest frequency of use (such as ceteareth 20 and aureth 7) usually contained less ethylene glycol units, working as surfactant-emulsifying agents.^62–64^ On the contrary, clinically used PEGylated nanomedicines typically contain PEG with the number of ethylene glycol units exceeding 40 to extend the circulation life-time of nanomedicines.^65^ Additionally, Tween-20 was used for the detection of anti-PEG antibodies in both human and animal sera in this study. Even if Tween-20 could compete with PEGylated materials for binding with antibodies against repeated ethylene glycol units, although there was currently no evidence to prove that, the lack of a significant difference in antibody affinity for MeO-PEG and OH-PEG suggested that anti-PEG antibodies in animals lacked selectivity for PEG terminal groups. On the contrary, the significant reduction in pre-existing anti-PEG antibody binding and complement activation in human serum when OH-PEG was used as the antigen compared with MeO-PEG suggested that there was at least a significant presence of anti-PEG antibodies targeting PEG termini in clinic.

We found the complement system in anti-PEG antibody positive individuals was in a hypersensitive state characterized by excessive activation of C3 and increased C5a level, which might serve as an accomplice to pre-existing anti-PEG antibodies, leading to enhanced macrophage uptake and poor stability, resulting in reduced circulation of nanomedicines modified with MeO-PEG, and even increased the risk for complement-associated allergy.^33^ It reminded us that it was necessary to check PEG antibody before the administration of PEGylated nanomedicines, or to try to use alternative products instead of MeO-PEG with lower binding affinity towards pre-existing anti-PEG antibodies, such as OH-PEG in medicine application.

## Materials and methods

### Cell lines, animals, and human serum

Human liver LO2 cells were provided by the Shanghai Institute of Cell Biology, Chinese Academy of Sciences (Shanghai, China). Human monocyte THP-1 cells were purchased from Dalian Meilun Biotechnology Co., Ltd. (Dalian, China). Male BALB/c and C57BL/6J mice, as well as SD rats (6-8 weeks old), were purchased from Shanghai SLAC Laboratory Animal Co., Ltd (Shanghai, China) and were kept under specific pathogen-free (SPF) conditions. All animal experiments were conducted in compliance with the guidelines approved by the Ethics Committee of Fudan University. Human serum samples were obtained from Jing’an District Central Hospital, Fudan University, Shanghai, China. The use of human samples in this study was approved by the Ethics Committee of Jing’an District Central Hospital (ethic approval no. 2022-23), and Fudan University School of Basic Medical Sciences (2021-C011).

### Reagents and antibodies

Ionizable cationic lipid SM-102 (8-[(2-hydroxyethyl)[6-oxo-6-(undecyloxy)hexyl]amino]-octanoic acid, 1-octylnonyl ester), DSPC (1,2-distearoyl-sn-glycero-3-phosphocholine), HSPC (Hydrogenated soy phosphatidylcholine), cholesterol, MeO-PEG2000-DSPE (1,2-distearoyl-sn-glycero-3-phosphoethanolamine-N-[methoxy(polyethylene glycol)-2000]), OH-PEG2000-DSPE (1,2-distearoyl-sn-glycero-3-phosphoethanolamine-N-[hydroxy(polyethylene glycol)-2000]), NH_2_-PEG2000-DSPE (1,2-distearoyl-sn-glycero-3-phosphoethanolamine-N-[amino(polyethylene glycol)-2000]), COOH-PEG2000-DSPE (1,2-distearoyl-sn-glycero-3-phosphoethanolamine-N-[carboxy(polyethylene glycol)-2000]), MeO-PEG2000-DMG (1,2-dimyristoyl-rac-glycero-3-[methoxy(polyethylene glycol)-2000]) and OH-PEG2000-DMG (1,2-dimyristoyl-rac-glycero-3-[hydroxy(polyethylene glycol)-2000]) were purchased from A.V.T. Pharmaceutical Co., Ltd. (Shanghai, China). MeO-PEG2000-PLGA45000 ([poly(L-lactide-co-glycolide)-45000]-[methoxy(polyethylene glycol)-2000]) and OH-PEG2000-PLGA45000 ([poly(L-lactide-co-glycolide)-45000]-[hydroxy(polyethylene glycol)-2000]) were purchased from Xi’an Ruixi Biological Technology Co., Ltd (Xi’an, China). MeO-PEG2000-PDLLA1750 ([poly(D,L-lactide)-1750]-[methoxy(polyethylene glycol)-2000]) and OH-PEG2000-PDLLA1750 ([poly(D,L-lactide)-1750]-[hydroxy(polyethylene glycol)-2000]) was purchased from Daigang Biomaterial Co., Ltd. (Jinan, China). Goat anti-rat IgM mu chain (HRP) (ab97180), goat anti-human IgM mu chain (HRP) (ab97205), goat anti-mouse IgM mu chain (HRP) (ab97230), goat anti-dog IgM H&L (HRP) (ab112835), goat anti-mouse IgG H&L (HRP) (ab97057), goat anti-human IgG Fc (HRP) (ab97225), goat anti-dog IgG H&L (HRP) (ab112852), recombinant anti-C3 antibody [EPR19394] (ab200999) and human complement C5a ELISA Kit (ab193695) were purchased from Abcam (Cambridge, MA). Goat anti-Rabbit IgG H&L (HRP), TMB (3,3’,5,5’-Tetramethylbenzidine) chromogen solution, gradient precast polyacrylamide gels (4–20%), Fast Silver Stain Kit and SDS-PAGE sample loading buffer (5×) were acquired from Beyotime Biotechnology (Nantong, China). Sheep red blood cell (SRBC, 4%) and hemolysin were acquired from Beijing Solarbio Science & Technology Co., Ltd. (Beijing, China). Hifair® _ 1st Strand cDNA Synthesis SuperMix for qPCR, Hieff® qPCR SYBR Green Master Mix and MolPure® Cell/Tissue Total RNA Kit were purchased from Yeasen Biotechnology Co., Ltd (Shanghai, China). Quant-iT™ RiboGreen™ RNA Assay Kit was purchased from Thermo Scientific Co., Ltd. (Rockford, IL, USA).

### Preparation and characterization of PEGylated nanocarriers

Lipid nanoparticles (LNP) were prepared using microfluidic mixing method. Briefly, a mixture of SM-102/DSPC/cholesterol/MeO-PEG-DMG or OH-PEG-DMG (molar ratio of 50:10:38.5:3.05) was dissolved in ethanol. mRNA was dissolved in a sodium acetate solution (25 mM, pH 5.0) and mixed with the lipid mixture at a volume ratio of 3:1 (aqueous: ethanol) to achieve a final molar ratio of 5.67:1 (ionizable N: P) through a microfluidic chip (Stainless steel, Aitesen, Suzhou, China). The mixture was immediately diluted tenfold with chilled PBS, and the external aqueous phase was replaced with PBS using Amicon® Ultra Centrifugal Filters (EMD Millipore, Billerica, MA, USA). The total RNA concentration and encapsulation efficacy were measured by RiboGreen assay. PEGylated liposomes (MeO-sLip and OH-sLip) were prepared using the thin film hydration and extrusion method, as reported in our previous papers. Briefly, a mixture of HSPC/cholesterol/mPEG-DSPE (molar ratio of 56.5:38.1:5.4) was dissolved in CHCl_3_ and vacuum-rotary evaporated to form a thin film, then dried overnight under vacuum to remove residual organic reagent. The lipid film was hydrated with normal saline at 60 °C. The lipid suspension was extruded through polycarbonate membranes with pore diameters of 200 nm, 100 nm and 50 nm to generate liposomes with an average particle size of 80 nm. PLGA nanoparticles were prepared using a single emulsion method. PLGA nanoparticles were prepared using a single emulsion method. Briefly, MeO-PEG2000-PLGA45000 or OH-PEG2000-PLGA45000 was dissolved in dichloromethane (10 mg/mL), and mixed with a 0.5% sodium cholate solution at a volume ratio of 4:1 (aqueous: ethanol). Subsequently, the mixture was subjected to ultrasonic emulsification for 8 minutes in an ice bath. After the removal of dichloromethane by rotary evaporation, uniform PLGA nanoparticles were collected by centrifugation at 3000 rpm to eliminate larger particles in the sediment, followed by centrifugation at 12000 rpm to remove smaller particles in the supernatant. Polymer micelles were prepared by thin film hydration method. MeO-PEG2000-PDLLA1750 or OH-PEG2000-PDLLA1750 were dissolved in ethanol and vacuum-rotary evaporated to form a thin film, then dried overnight under vacuum. The dried lipid film was hydrated with saline at 37°C to form micelles self-assembly.

### Evaluation of detergent capability of CHAPS and Tween-20

Medium-binding 96-well ELISA plates were coated with 2 µg of MeO-PEG-DSPE per well. After rinses with 0.05%Tween-20/PBS or 0.05%CHAPS/PBS, wells were blocked with 5% BSA-PBS at 37 °C for 1 h. Purified human IgM or IgG (800 ng/mL, non-anti-PEG type, purchased from Bethyl Laboratories) were added to different concentration of anti-PEG IgM (cAGP4) or IgG (c3.3) as an indicator of nonspecific binding. The mixture was incubated in the wells at 37 °C for 1 h. After rinsed with 0.05%Tween-20/PBS or 0.05%CHAPS/PBS, HRP-labeled goat anti-IgM or IgG antibody was added to bind with IgM or IgG at 37 °C for 1 h. TMB substrate was added for 8 min to visualize HRP, and the reaction was terminated using 0.18 M sulfuric acid. UV absorbance at 450 nm was then measured.

The blank medium-binding 96-well ELISA plates were added with goat anti human IgM or IgG (HRP) and incubated at 37_ for 1 h, PBS was used as the control group. After rinsed with 0.05%Tween-20/PBS or 0.05%CHAPS/PBS, TMB substrate was added for 8 min to visualize HRP, and the reaction was terminated using 0.18 M sulfuric acid. UV absorbance at 450 nm was then measured.

### Measurement of PEG antibodies

ELISA was utilized to evaluate the binding affinity between anti-PEG antibody and PEG. Medium-binding 96-well ELISA plates were coated with 2 µg of PEGylated materials (such as MeO-PEG-DSPE, OH-PEG-DSPE, MeO-PEG-DMG, etc.) or 20 µg of PEGylated nanocarriers (such as sLip, LNP, etc.) per well. After rinses with 0.05% PBST thrice, wells were blocked with 5% BSA-PBS at 37 °C for 1 h. Serum was gradually diluted with 1% BSA-PBS and incubated in the microtiter wells at 37 °C for 1 h. Following rinses with 0.05% PBST, HRP-labeled goat anti-IgM or IgG antibody was added to bind with IgM or IgG at 37 °C for 1 h. TMB substrate was added for 8 min to visualize HRP, and the reaction was terminated using 0.18 M sulfuric acid. UV absorbance at 450 nm was then measured.

### Isothermal titration calorimetry

To minimize heat of dilution release from solvent dilution due to the high concentration of salt ions and proteins in the serum, we added blank rat serum (which does not contain anti-PEG antibodies negative) to PEG materials-DSPE to a final concentration of 20%. Approximately 300 μL of 20% serum sample was carefully injected into the sample cell of MicroCal PEAQ-ITC (Malvern Instruments, Malvern, UK), and 60 μL of blank serum mixture was loaded to the syringe for titration into the serum sample. Experimental parameters were set as follows: running temperature 25°C, paddle speed 750 rpm, total drop 19, drop interval 60 s, initial delay 60 s. The volume of each drop was 2 μL, except for the first drop (0.4 μL). A 20% blank rat serum was used as the reference, with a reference power set to 10 μcal/s. Since there is no standard for pre-existing anti-PEG antibodies at present, the concentration of anti-PEG antibodies in the human serum was unknown, therefore, the sample concentration is set based on the total antibody IgM and IgG concentration, approximately 20 μM; the concentration of PEG-DSPE was 200 μM.

### The complement activation of LNP in serum

BALB/c mice (5 mg HSPC/kg), SD rats (2.5 mg HSPC/kg) or beagle dogs (1 mg HSPC/kg) were intravenously injected with empty PEGylated liposomes to generate anti-PEG antibodies. After 6 days, blood was sampled and the serum was separated. The complement activation of LNP in serum was assessed by the detection of C3 cleavage through Western blot assay. LNP were mixed with either pre-existing anti-PEG antibodies positive/negative human serum or sLip-stimulated/blank mouse/rat/dog serum at a final lipid concentration of 1 mg/mL and incubated at 37 °C for 1 h. The mixture was then diluted 200 times with 10 mM EDTA-PBS to terminate complement activation. C5a level in the mixture was measured via human C5a ELISA kit (abcam 193695) according to the manufacturer’s instructions. For C3 cleavage, the mixture was mixed with 5× loading buffer. After heating for 10 minutes at 95°C, the sample was loaded onto a 4%–20% gradient polyacrylamide gel for electrophoresis and transferred to a PVDF membrane. Nonspecific binding sites on the PVDF membrane were blocked by incubating in PBST containing 5% BSA at room temperature for 2 h, followed by overnight incubation at 4°C with recombinant Anti-C3 antibody (1:1000). After six washes with PBST (5 min each) to remove residual primary antibody, HRP-labeled Goat anti-rabbit IgG H&L (1:1000) was added as the second antibody. The signals of C3 and cleavage products were imaged using a ChemiScope 6000 (Clinx Co., Ltd), and the data were analyzed with Image J software.

### Characterization of protein corona of LNP

To compare the *in vitro* IgM/IgG adsorption abilities of MeO-LNP and OH-LNP, LNP were incubated with pre-existing anti-PEG antibodies positive/negative human serum at a final lipid concentration of 0.5 mg/mL at 37 °C for 1 h. The mixtures were diluted with chilled PBS to reduce serum viscosity and density, then centrifuged (14000 × g, 30 min) and rinsed with chilled PBS (300 μL) thrice to collect protein coronas formed on the surface of the LNP. Protein aliquots were separated using a 4%–20% gradient polyacrylamide gel and stained with the Fast Silver Stain Kit or transferred to a PVDF membrane. Human IgM and IgG were detected using the appropriate secondary HRP-conjugated antibody.

### Cell culture and THP-1 derived macrophage differentiation

Human liver L02 cells were cultured in high-glucose DMEM containing 4.5g/L glucose, 10%FBS, 100 units/mL penicillin and 100 μg/mL streptomycin with 5% CO2 at 37 °C. human monocytic cell line THP-1 were cultured in calcium-free RPMI 1640 containing 10%FBS, 100 units/mL penicillin, 100 μg/mL streptomycin and 0.05 mM β-mercaptoethanol with 5% CO2 at 37 °C. THP-1 cells were seeded in 24-well plates at a density of 200,000 cells per well. Phorbol-12-myristate-13-acetate (PMA, 100 ng/mL) was added in culture medium to induce the differentiation of THP-1 into macrophages. After 2 days, the medium was changed to complete culture medium without PMA to rest the cells for 3 days before the experiment.

### Cellular uptake of LNP by human hepatocytes and macrophages

Human liver L02 cells were seeded at a density of 60,000 cells per well in 24-well plates and allowed to grow for 16 hours to achieve sufficient adhesion. Human macrophages were derived from THP-1 cells through treatment with PMA, following the method mentioned above. LNP encapsulating eGFP mRNA were pre-incubated with an equal volume of pre-existing anti-PEG antibodies positive/negative human serum at 37°C for 1 h, and then incubated with cells in pure DMEM (for L02) or RPMI 1640 (for macrophages) medium at an RNA concentration of 2 μg/mL for 2 h at 37°C. After washing thrice with chilled PBS, cells were harvested, and total RNA was extracted using the MolPure® Cell/Tissue Total RNA Kit. Subsequently, 1 μg RNA was reverse transcribed into complementary DNA (cDNA) using the Hifair® _ 1st Strand cDNA Synthesis SuperMix according to the manufacturer’s instructions. The relative levels of eGFP mRNA were analyzed via RT-qPCR on a LightCycler 480. Fold changes were calculated using the ΔΔCt method, with human GAPDH as the reference gene for normalization. The primers used for validation are listed in the Supplementary Table 3.

### Stability of mRNA-LNP in human serum

LNP encapsulating eGFP mRNA (50 μg mRNA/mL) were incubated with a 4-times volume of pre-existing anti-PEG antibodies positive/negative human serum in the presence of RNase at 37°C for 2 h to degrade any leaked mRNA from the LNP. PBS and 1% Triton X-100 were employed as negative and positive controls, respectively. After 2 h, Diethyl pyrocarbonate (DEPC) was added to inhibit the activity of RNase, and L02 cells (100,000 cells per sample) were incorporated to introduced GAPDH mRNA as an internal standard. Total RNA was extracted from the serum-cell mixtures, and the relative levels of eGFP mRNA remaining in LNP were analyzed using the same methods as aforementioned.

### The immunogenicity of LNP

C57BL6 mice (40 mg lipid per kg of mice) or Beagle dogs (60 mg lipid per kg of dogs) were intravenously injected three doses (by weekly) of LNP. Blood was sampled 7 days after each injection and kept at room temperature for 30 min. Serum was collected after centrifugation at 1000 × g for 10 min and stored at −80 °C. Anti-PEG IgM/IgG titrations in serum were detected by ELISA assay using MeO-PEG-DMG (MeO-LNP) or OH-PEG-DMG (OH-LNP) as antigens.

### Statistical analysis

Data are means ± standard deviations (SDs) and analyzed by Student’s t-test, one-way ANOVA or two-way ANOVA followed their respective applicable principles with GraphPad Prism 9.0. *p < 0.05* was considered statistically significant (n.s.: non-significance *p > 0.05*, *0.01 <* ∗*p < 0.05*, *0.001 <* ∗∗*p < 0.01*, ∗∗∗*p < 0.001*).

## Data availability statement

All data that support the findings of this study are available within the main text or the Supporting information. Other data are available from the authors upon reasonable request.

## Acknowledgement

The authors declare that there are no known competing financial interests or personal relationships that could have appeared to influence this paper. This work was supported by the National Natural Science Fund of China (82125035, 32330058, 82273866, 82361168639), Shanghai Education Commission Major Project (2021-01-07-00-07-E00081) and China Postdoctoral Science Foundation (GZC20240305).

## Supporting information

The Supporting information is available free of charge online.

**Supplemental table 1-3:** Fitting parameters from ITC analysis; Characterization of lipid nanoparticles; Primer sequences

**Supplemental Figure 1-21:** The cross-reactivity test of secondary antibody; Evaluation of detergent capability of CHAPS and Tween-20; The inhibitory experiment; The relationship between OD_450nm_ value and titration of IgG; The prevalence of anti-PEG antibodies in human; The influence of Tween-20 on the binding of antibodies; PEG terminus selectivity of antibodies; The binding of antibodies with other PEG materials; ITC data; The binding of antibodies with other PEGylated nanocarriers; C5a level in human serum after incubation with LNPs; The effect of complement deactivation on biological functions of antibodies; The cleavage of C3 protein in human serum; C5a level in untreated human serum; The binding affinities of antibody in rats with PEG materials; The immunogenicity of LNP in mice; The uncropped and unprocessed gel

