## Supplemental information for "Hydroxy polyethylene glycol: a solution to evade human pre-existing anti-PEG antibodies for efficient delivery"

Supplemental Table

Supplemental Table 1. Fitting parameters ΔH and K_D_ from ITC analysis of the integrated heats with independent-site model. For OH-PEG, K_D_ and ΔH was not fitted due to the lack of specific binding. ΔH was the heat difference between the first and last injection, and K_D_ was marked as ND (Not Detectable).

| Donor number | MeO-PEG | | OH-PEG | |
| --- | --- | --- | --- | --- |
|  | ΔH (kcal/mol) | K_D_ (M) | ΔH (kcal/mol) | K_D_ (M) |
| #670 | -9.5±1.3 | (5.6±1.8)e-6 | 1.5 | ND |
| #1740 | -18.4±6.4 | (6.2±3.0)e-6 | 1.8 | ND |
| #886 | -12.7±1.6 | (3.6±1.2)e-6 | 0.6 | ND |
| #908 | -8.1±1.1 | (3.5±1.4)e-6 | 1.3 | ND |
| #638 | -7.7±0.7 | (4±0.8)e-6 | 1.1 | ND |
| #640 | -20.0±9.0 | (14.1±7.3)e-6 | 1.0 | ND |

Supplemental Table 2. Characterization of lipid nanoparticles. The percentage indicates the degree of modification of PEG-DMG in LNP. Data are means ± SDs (n = 3).

|  | 1.5%MeO-LNP | 3%MeO-LNP | 5%MeO-LNP | 1.5%OH-LNP | 3%OH-LNP | 5%OH-LNP |
| --- | --- | --- | --- | --- | --- | --- |
| Size (nm) | 75.4±1.3 | 65.9±0.4 | 63.5±1.7 | 75.3±0.5 | 72.7±0.4 | 67.9±0.7 |
| PDI | 0.08±0.01 | 0.09±0.01 | 0.08±0.01 | 0.08±0.02 | 0.09±0.02 | 0.09±0.01 |

Supplemental Table 3. Primer sequences for qPCR.

| **Primers** | **Primer sequences (5′-3′)** |
| --- | --- |
| **eGFP-F** | AAGGACGACGGCAACTACAA |
| **eGFP-R** | CGATGTTGTGGCGGATCTTG |
| **Human GAPDH-F** | GTCTCCTCTGACTTCAACAGCG |
| **Human GAPDH-R** | ACCACCCTGTTGCTGTAGCCAA |

Supplemental Figures

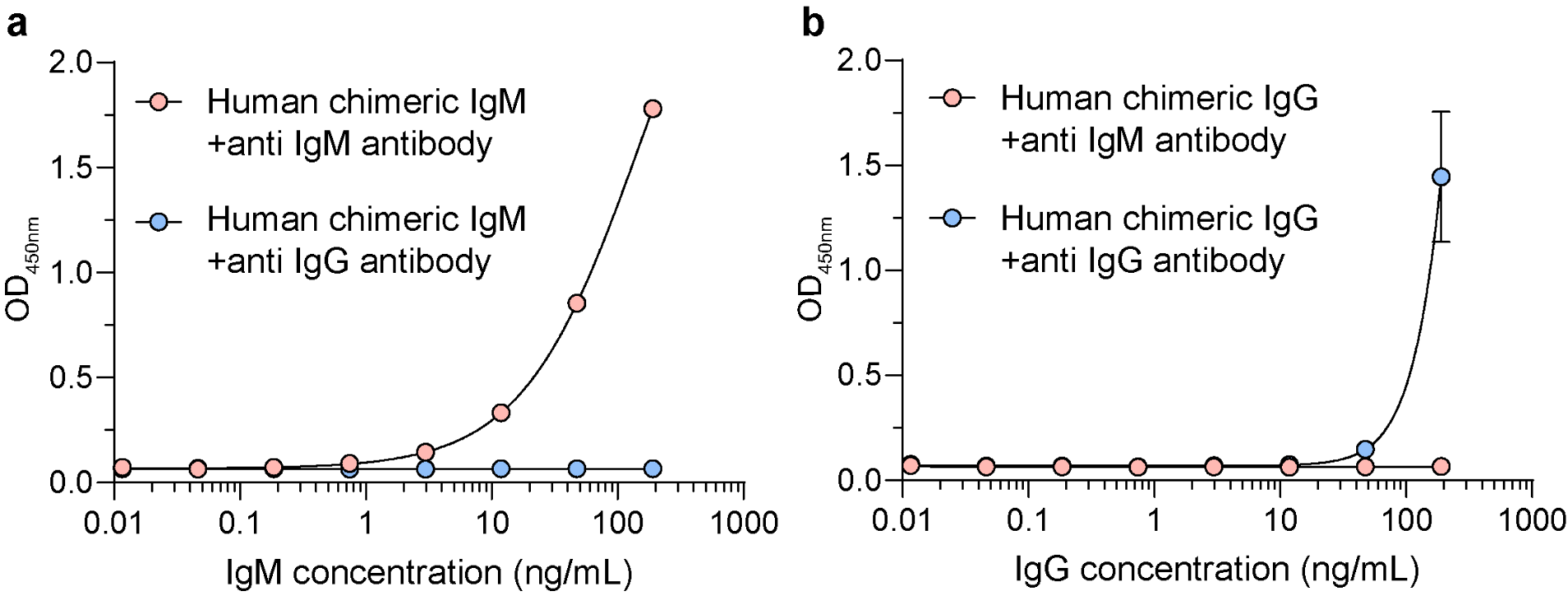

**Supplemental Figure 1**. **The cross-reactivity test of the secondary antibody.** The binding curve of human chimeric anti-PEG IgM **(a)** or IgG **(b)** with MeO-PEG using HRP goat anti human IgM or IgG as secondary antibodies. Data are means ± SDs (n = 3).

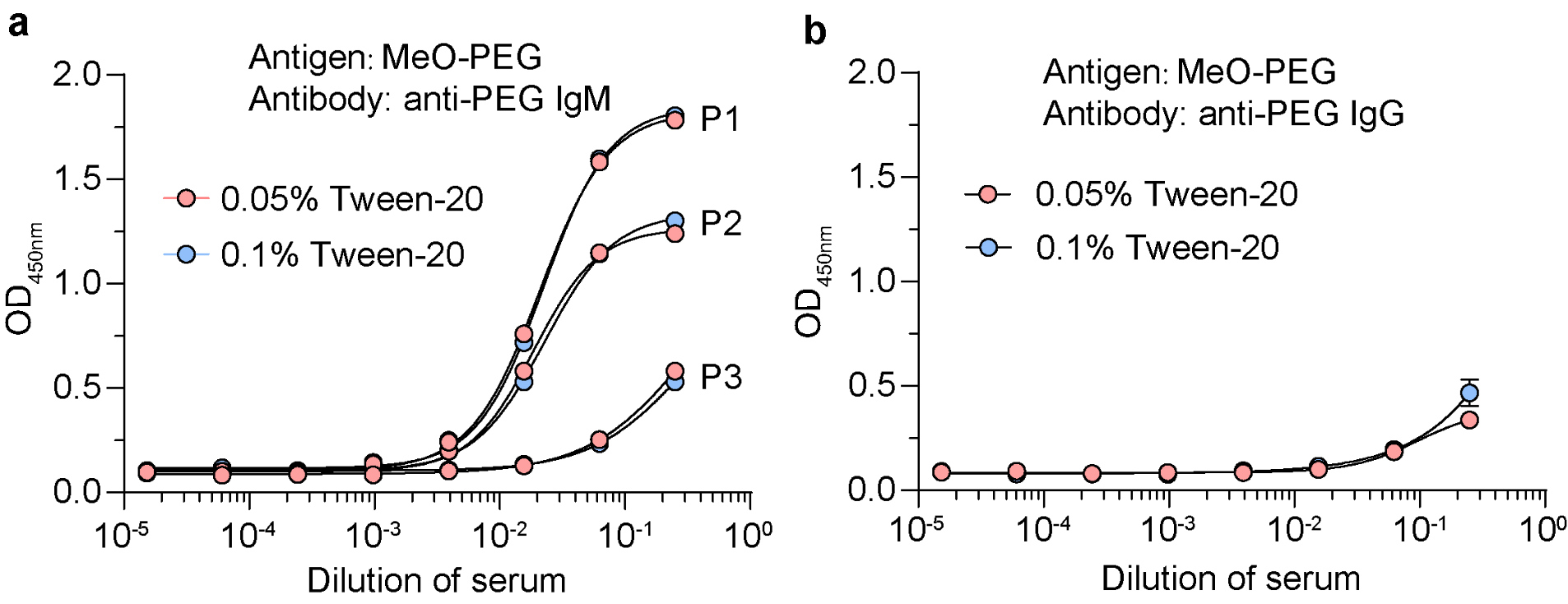

**Supplemental Figure 2**. **The binding of pre-existing anti-PEG antibodies with MeO-PEG using different concentrations of Tween-20 as wash buffer**. The binding curve of pre-existing anti-PEG IgM **(a)** or IgG **(b)** with MeO-PEG-DSPE. P1, P2 and P3 represented the binding curve of three distinct human samples. Data in (b) are shown as means ± SDs using the same samples in (a) (n = 3).

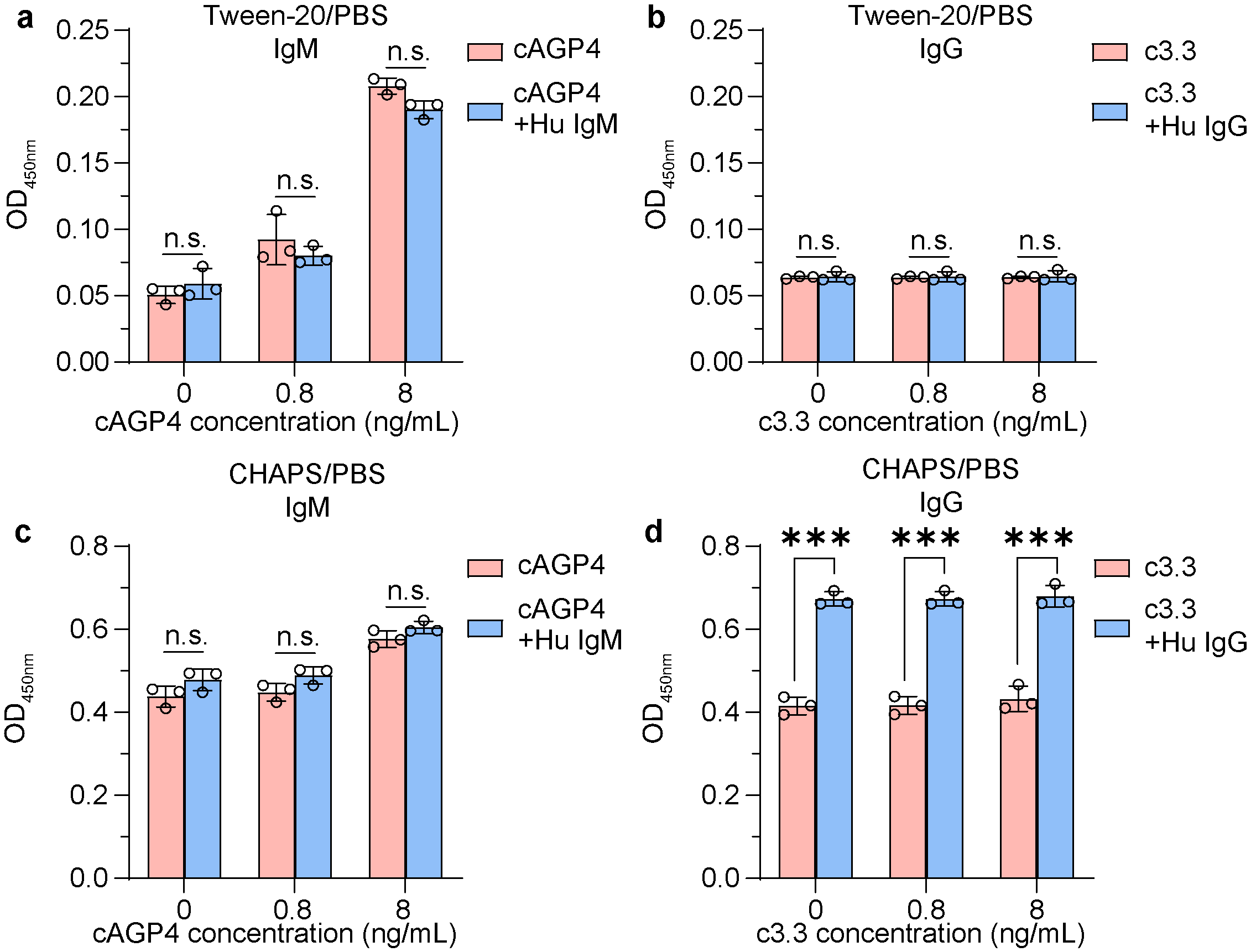

**Supplemental Figure 3. The binding of anti-PEG antibodies with MeO-PEG in the presence or absence of human standard antibodies.** The binding of anti-PEG IgM (a) or IgG (b) with MeO-PEG-DSPE using 0.05% Tween-20/PBS as washing buffer. The binding of anti-PEG IgM (c) or IgG (d) with MeO-PEG-DSPE using 0.05% CHAPS/PBS as washing buffer. Data are means ± SDs (n = 3). Statistical significance is evaluated using Two-way ANOVA with GraphPad Prism 9.0 (n.s. indicates non-significant, ***p<0.001).

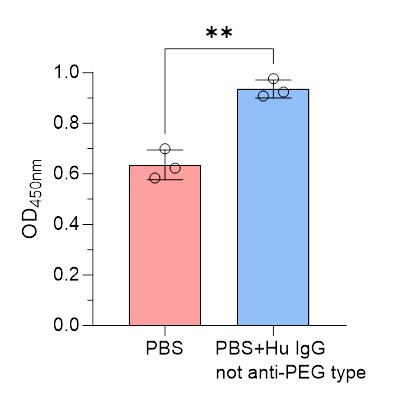

**Supplemental Figure 4. Washing capabilities of 0.1%CHAPS/PBS on non-specific primary antibodies.** Blank ELISA plants coated with MeO-PEG-DSPE were added with PBS or human standard IgG, and rinsed with 0.1%CHAPS/PBS. Data are means ± SDs (n = 3). Statistical significance is evaluated using Student t test with GraphPad Prism 9.0 (**p<0.01).

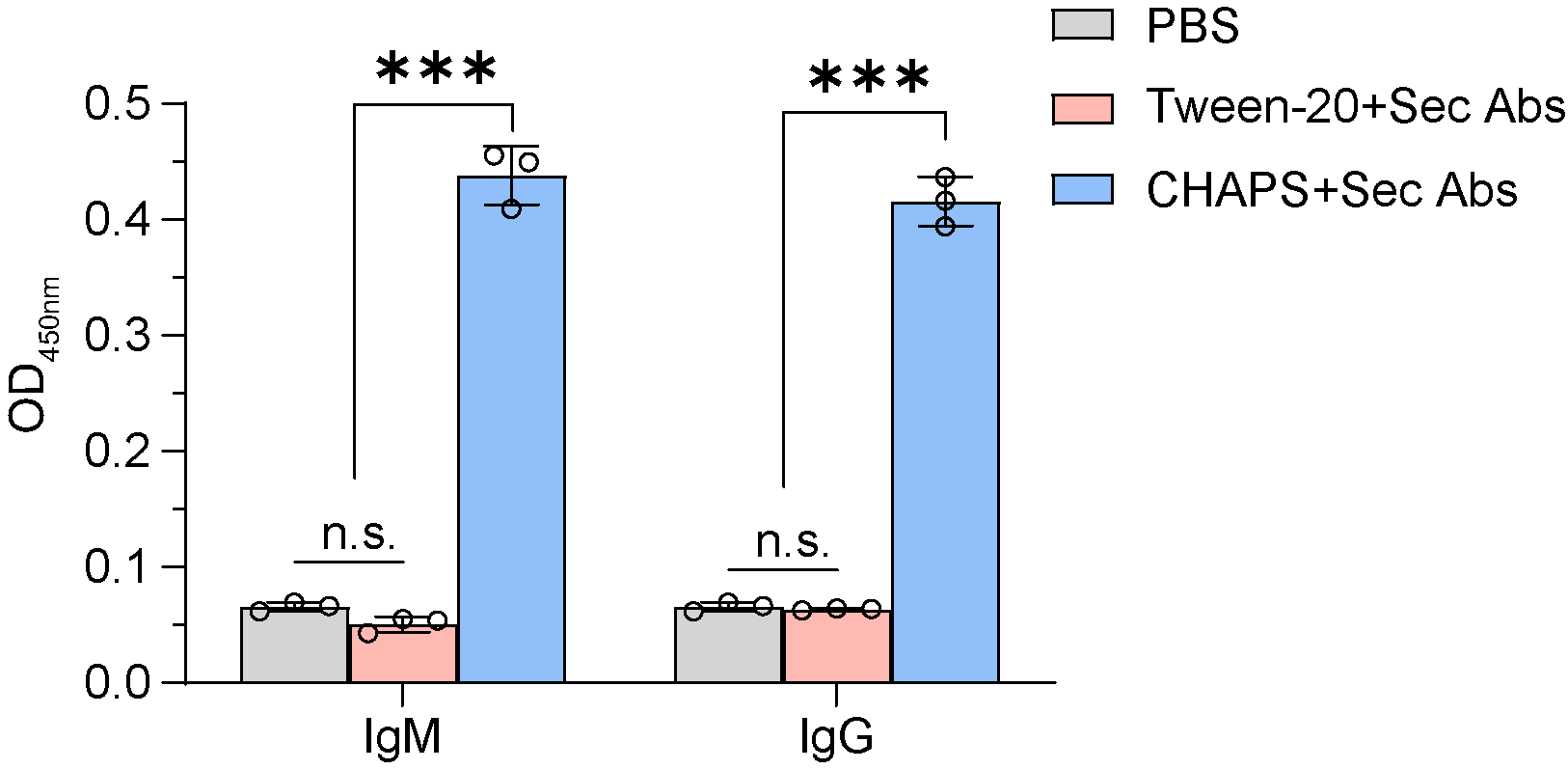

**Supplemental Figure 5. Washing capabilities of different detergents on non-specific secondary antibodies**. Blank ELISA plants were added with HRP labeled secondary antibodies or PBS, and rinsed with Tween-20 or CHAPS. Data are means ± SDs (n = 3). Statistical significance is evaluated using Two-way ANOVA with GraphPad Prism 9.0 (n.s. indicates non-significant, ***p<0.001).

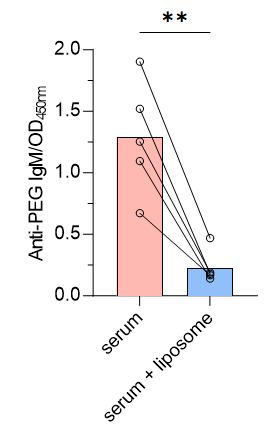

**Supplemental Figure 6**. **The inhibitory experiment of pre-existing anti-PEG IgM with PEGylated liposomes as inhibitors.** Levels of anti-PEG IgM before and after adding the same volume of empty PEGylated liposomes (final PEG content is 30 μg/well). Data are shown in dots and lines (n = 5) and statistical significance is evaluated using paired Student's t-test with GraphPad Prism 9.0 (**p<0.01).

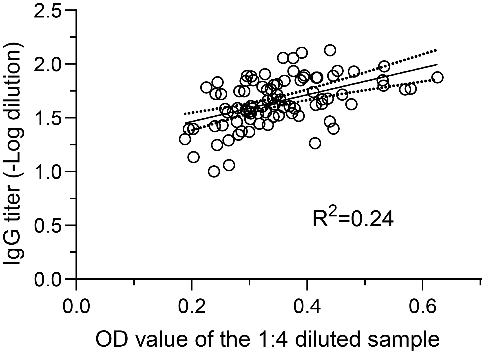

**Supplemental Figure 7. The relationship between single point OD_450nm_ value (1:4 diluted) and titration of IgG (n=90).** Titration was calculated as -log(dilution, whose OD_450nm_ was 2.1 times of the control buffer) using the embedded equation of specific binding in GraphPad Prism 9.0. Linear regression line (Y=1.249*X+1.215, R^2^=0.24) was shown in solid line. Dotted lines showed 95% prediction intervals.

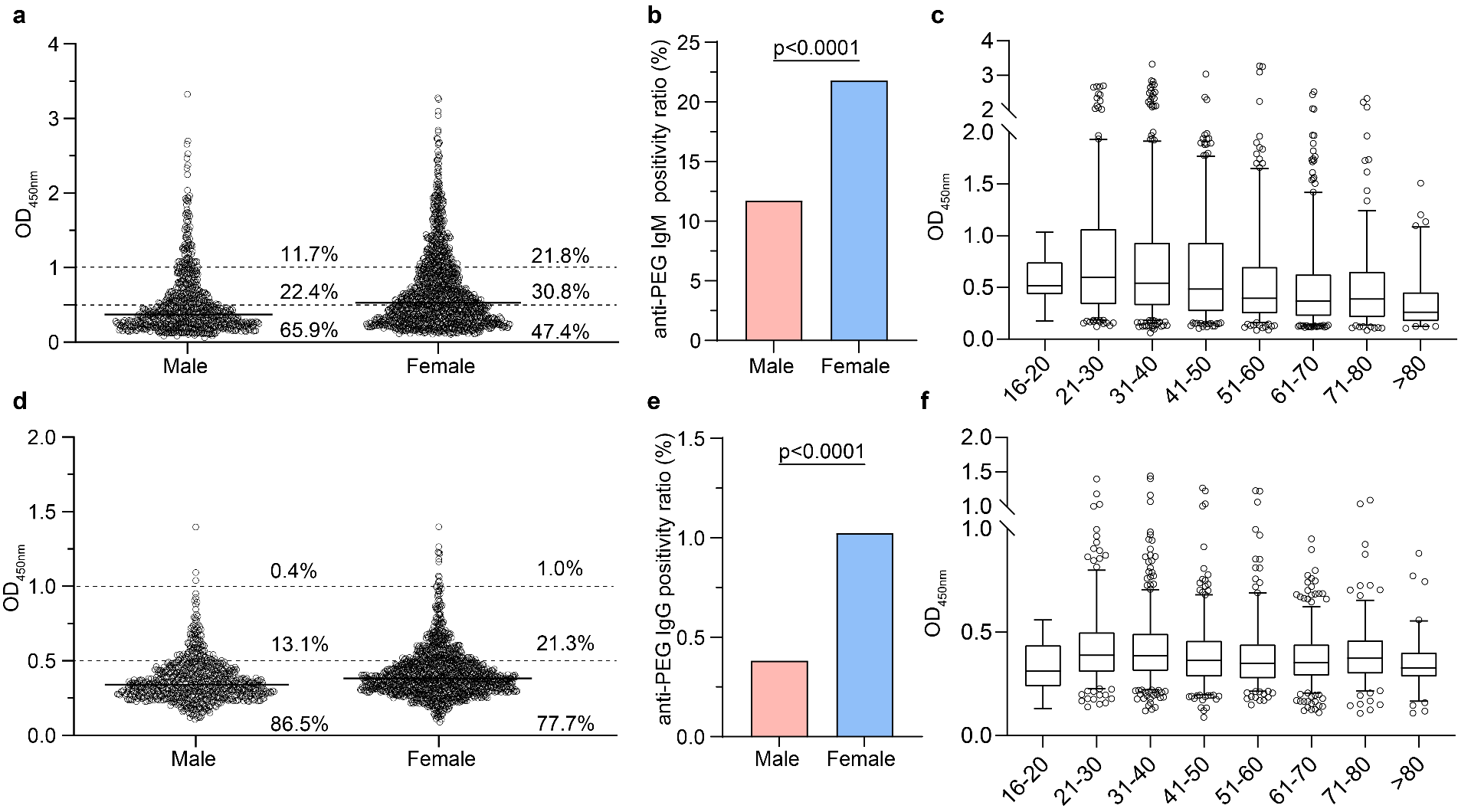

**Supplemental Figure 8. The prevalence of anti-PEG antibodies in human.** Gender distribution of pre-existing anti-PEG IgM **(a)** and IgG **(d)** in total population (male 784 and female 1290). Gender differences in positivity ratios of anti-PEG IgM (males 92 of 784 and females 281 of 1290) **(b)** and IgG (males 3 of 784 and females 13 of 1290) **(e)**. OD_450nm_ values (1:4 diluted) of anti-PEG IgM **(c)** and IgG **(f)** is shown versus donor age (each age groups with n ≥ 18). Data are means±SDs and analyzed with GraphPad Prism 9.0.

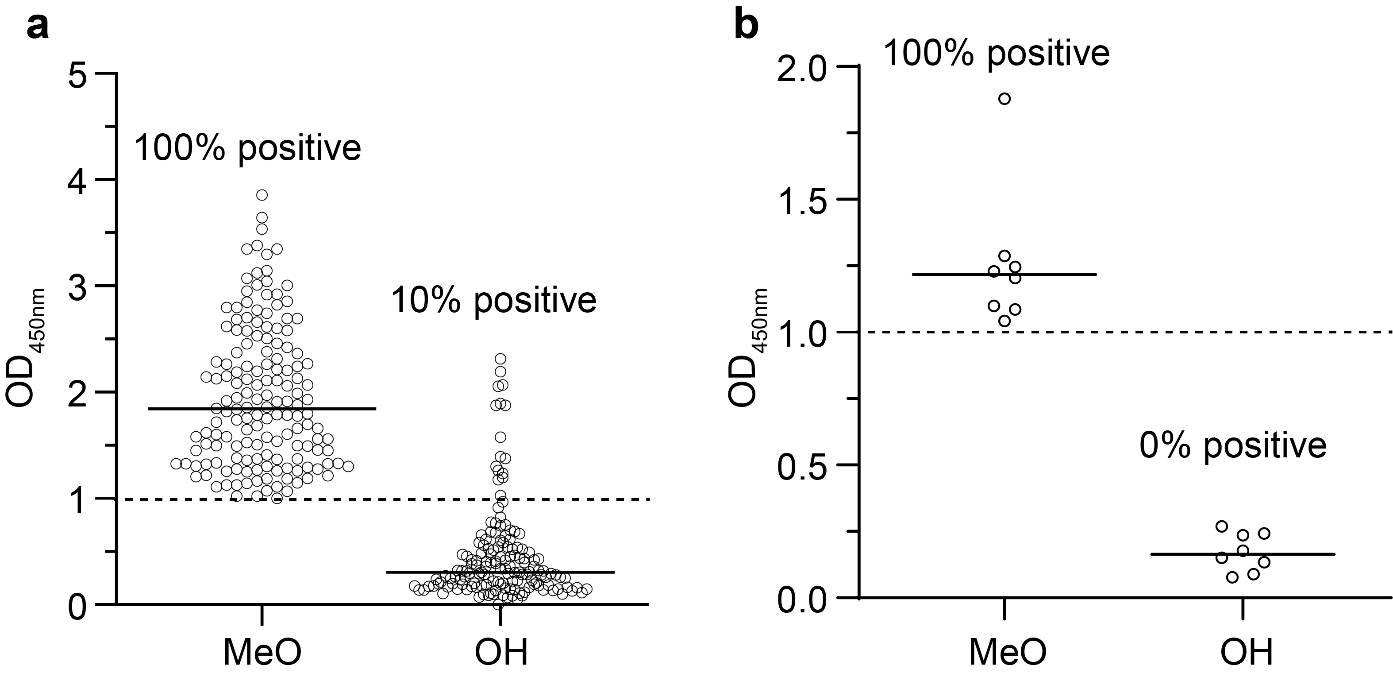

**Supplemental Figure 9. PEG terminus selectivity of human pre-existing anti-PEG antibodies in samples from Pudong Hospital cohort (Shanghai, China).** The positive rate of anti-PEG IgM **(b)** or IgG **(e)** in strongly positive human samples against OH-PEG-DSPE (n=155 and 8). Data are shown in dots and means, and analyzed with GraphPad Prism 9.0.

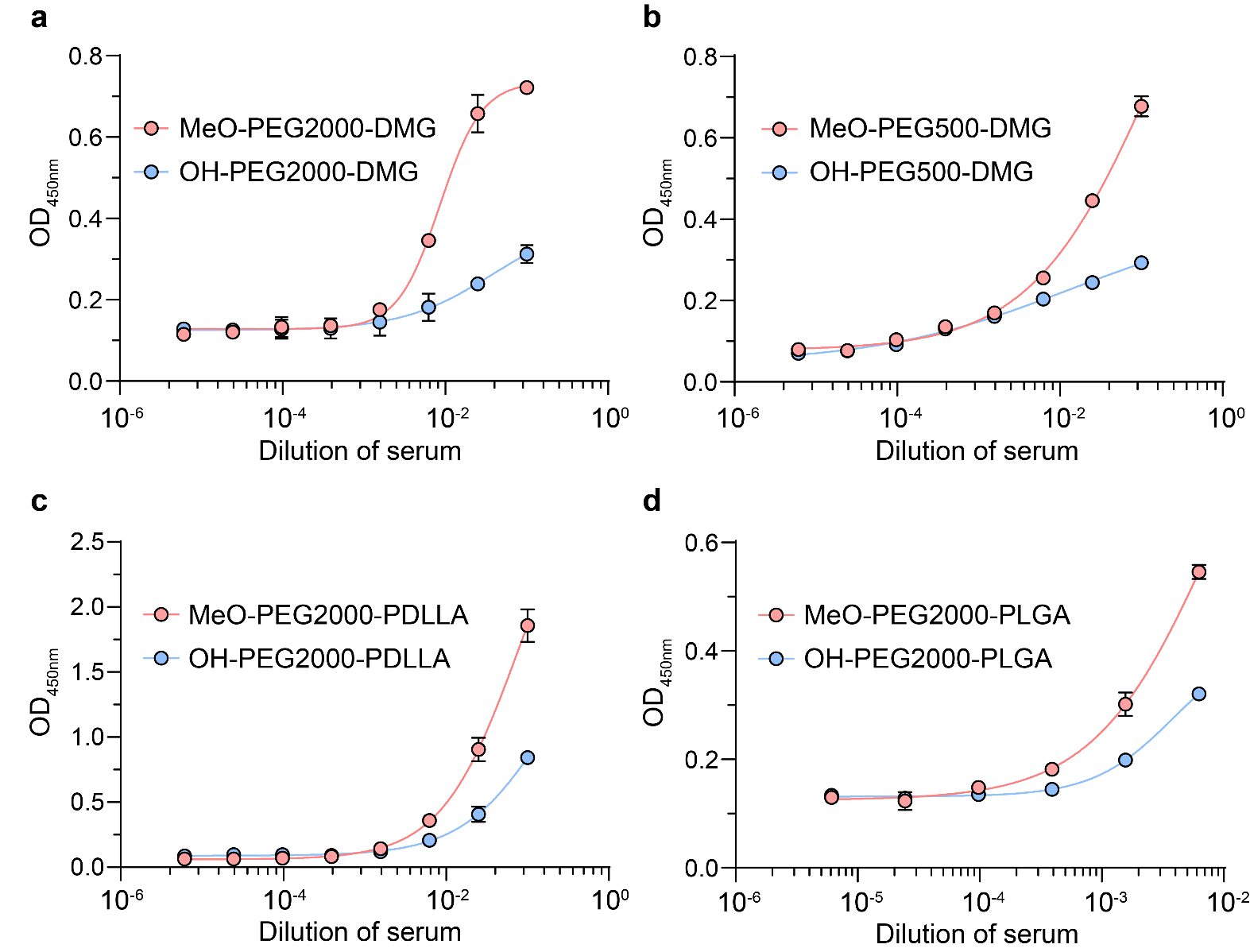

**Supplemental Figure 10. The binding of pre-existing anti-PEG antibodies with PEG materials of different molecular weights and hydrophobic groups.** The binding curve of pre-existing anti-PEG IgM with PEG2000-DMG **(a)**, PEG500-DMG **(b)**, PEG2000-PDLLA **(c)** and PEG2000-PLGA **(d)** with methoxy or hydroxy as the terminal group. Data are means ± SDs (n = 3).

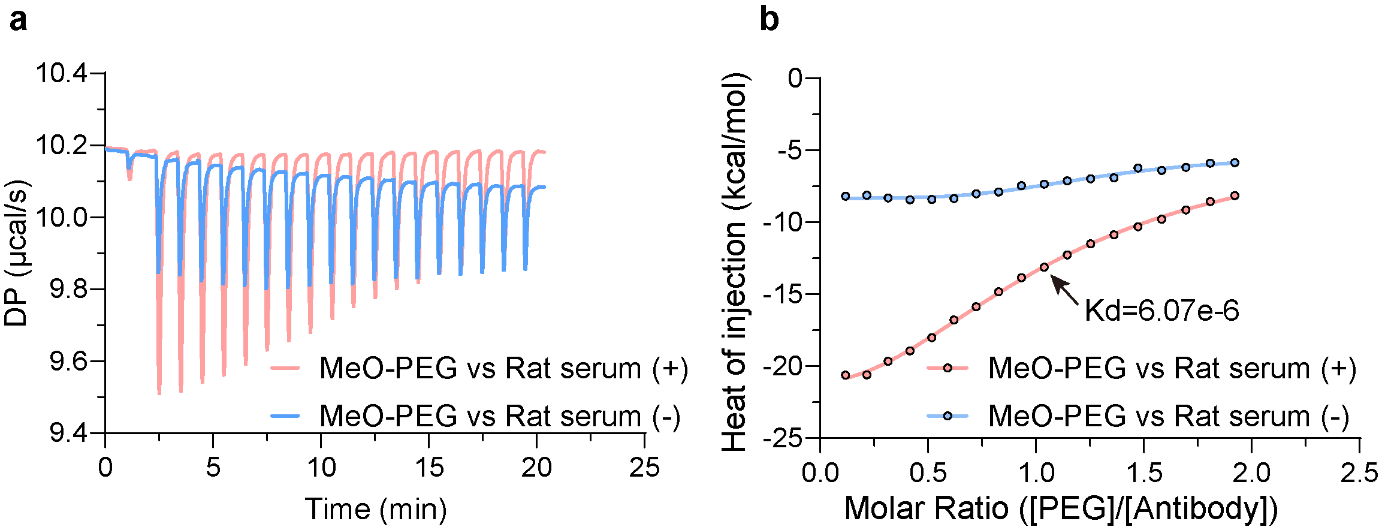

**Supplemental Figure 11. Typical data obtained from isothermal titration calorimetry measurements of serum titrated with MeO-PEG.** Real-time thermogram (a) and the area of each peak (b) represent the heat release during titration of rat serum with MeO-PEG.

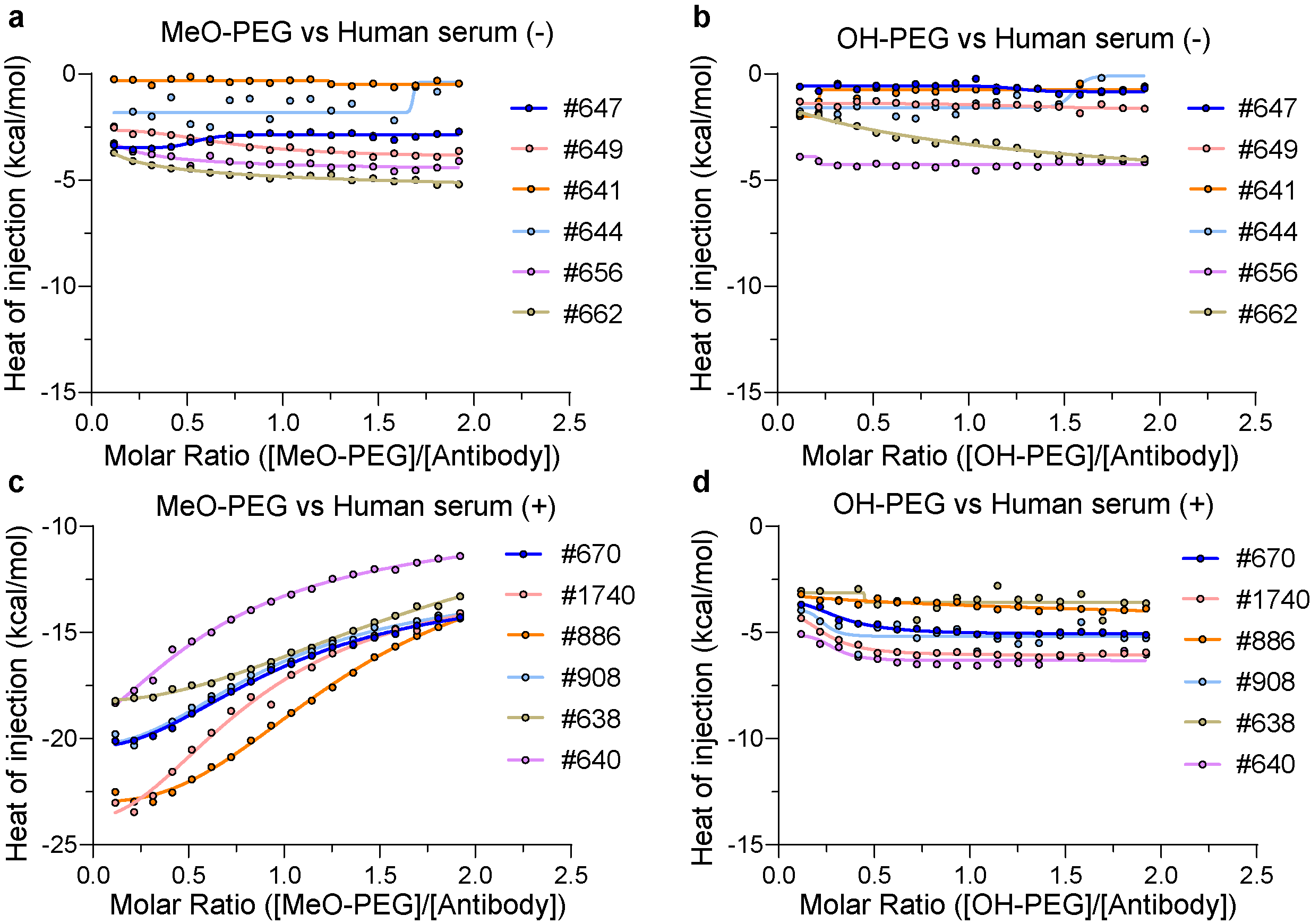

**Supplemental Figure 12. The heat release during titration of human serum with PEG.** Anti-PEG antibodies negative human serum (OD<0.5 in ELISA) titrated with MeO-PEG **(a)** or OH-PEG **(b)**. Anti-PEG antibodies positive human serum (OD>1 in ELISA) titrated with MeO-PEG **(c)** or OH-PEG **(d)**.

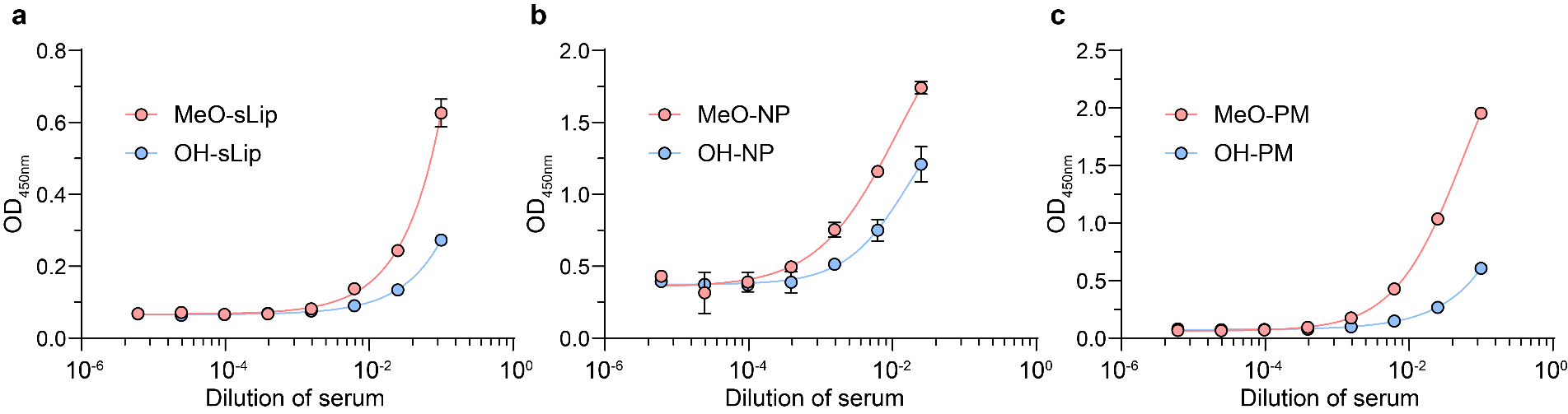

**Supplemental Figure 13. The binding of pre-existing anti-PEG antibodies with other PEGylated nanocarriers.** The binding curve of pre-existing anti-PEG IgM with liposomes **(a)**, PLGA nanoparticles **(b)** and polymer micelles **(c)** with MeO-PEG or OH-PEG modification. MeO-sLip, MeO-NP and MeO-PM represents liposomes, PLGA nanoparticles and polymer micelles modified with MeO-PEG. OH-sLip, OH-NP and OH-PM represents liposomes, PLGA nanoparticles and polymer micelles modified with OH-PEG. Data are means ± SDs (n = 3).

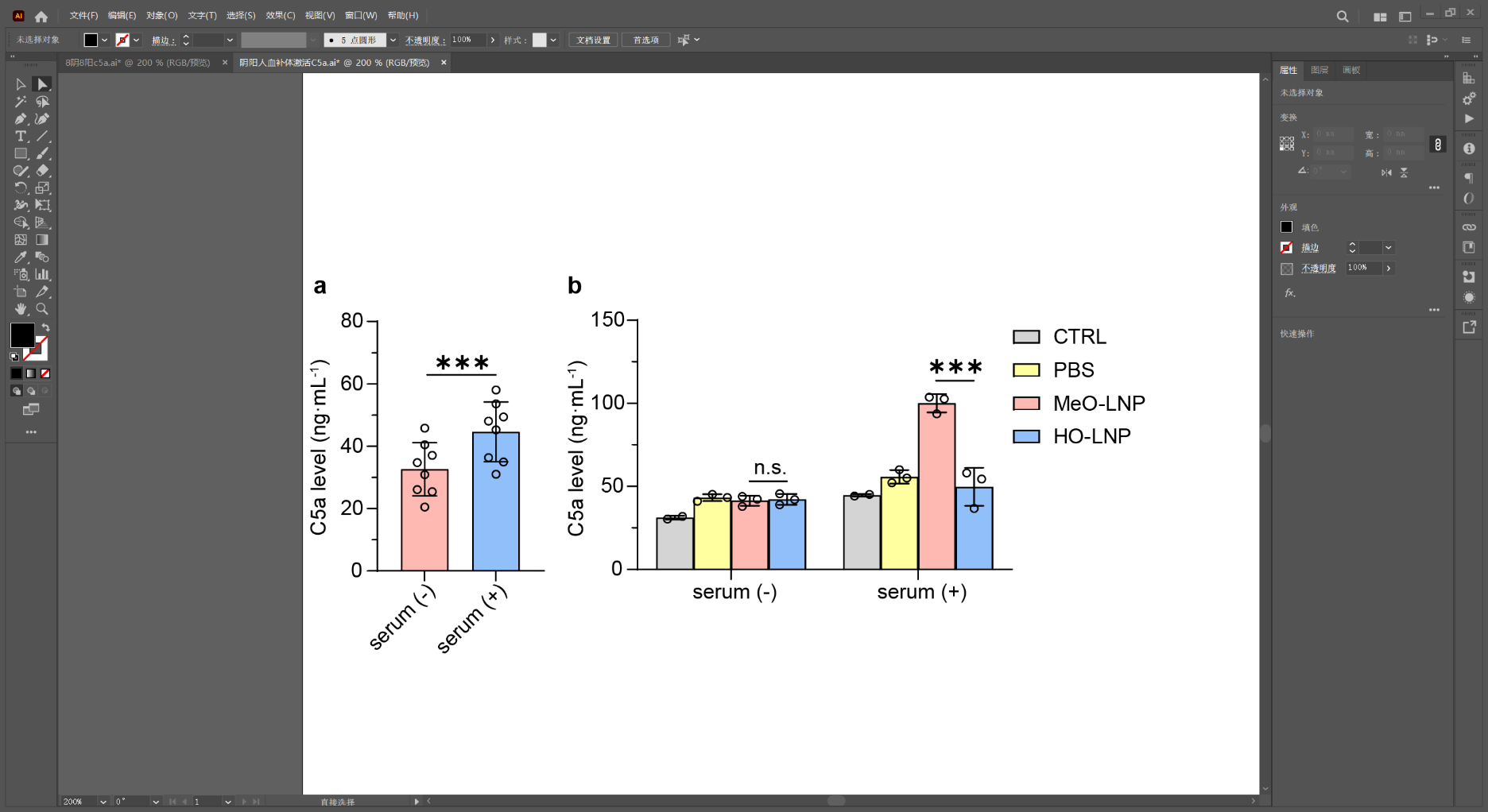

**Supplemental Figure 14. C5a level in human serum after incubation with LNPs.** ELISA detection of C5a levels in pre-existing anti-PEG antibody negative serum (serum(-)) or positive serum (serum(+)) after incubation with LNP for 1 h. Data are means ± SDs (n = 3). Statistical significance is evaluated using Two-way ANOVA with GraphPad Prism 9.0 (n.s. indicates non-significant, *^***^p<0.001*).

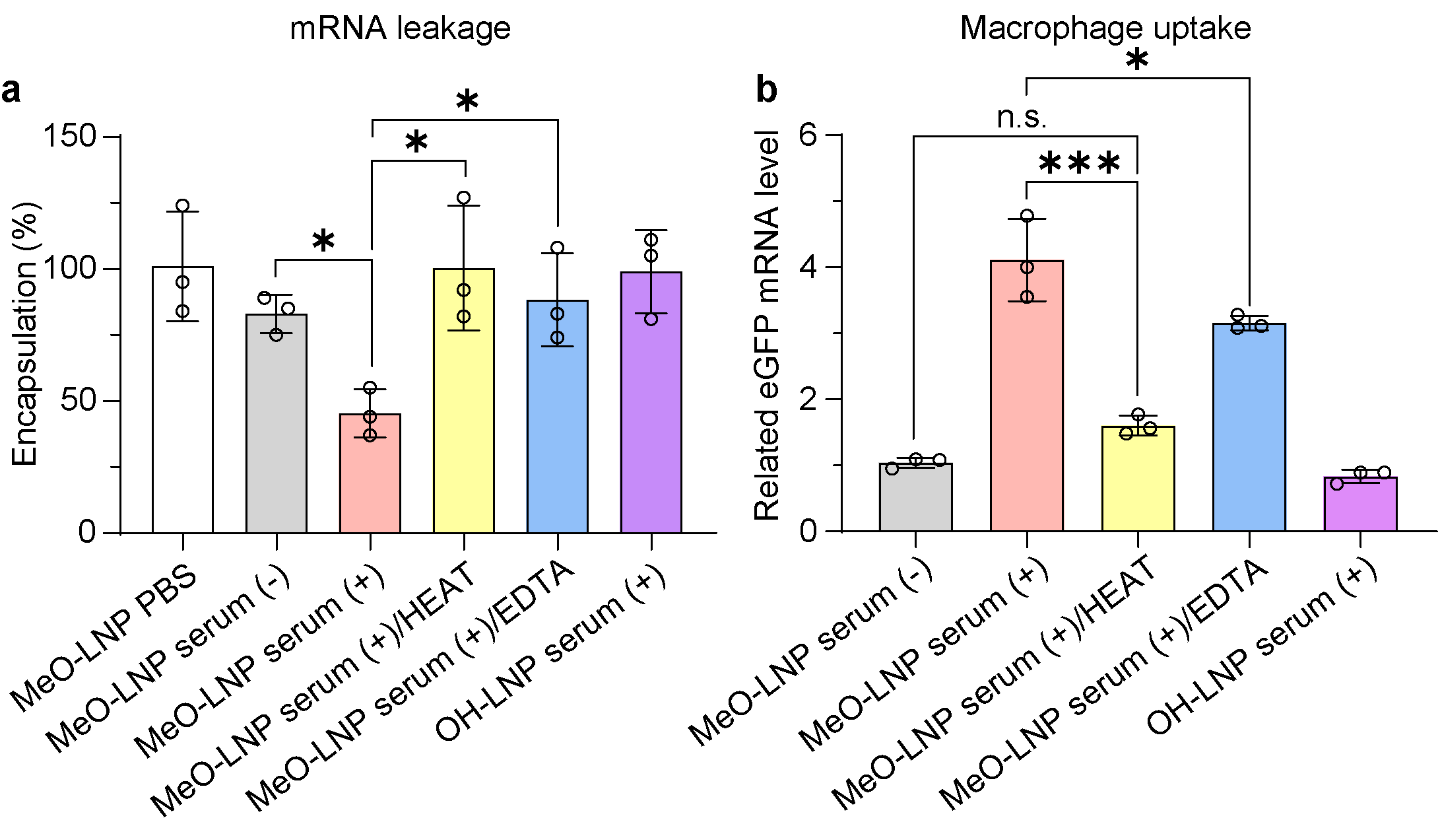

**Supplemental Figure 15. The effect of complement deactivation on the biological functions of human pre-existing antibodies on LNP. (a)** The remaining eGFP mRNA encapsulated in LNP after incubation with human serum with different pre-treatment for 2 h in the presence of RNase. **(b)** Uptake of eGFP-LNP by THP-1 derived macrophage in the presence of human serum with different pre-treatment, quantified by RT-qPCR. Data are means±SDs (n = 3). Statistical significance is evaluated using One-way ANOVA with GraphPad Prism 9.0 (n.s. indicates non-significant, *^*^p<0.05, ^***^p<0.001*).

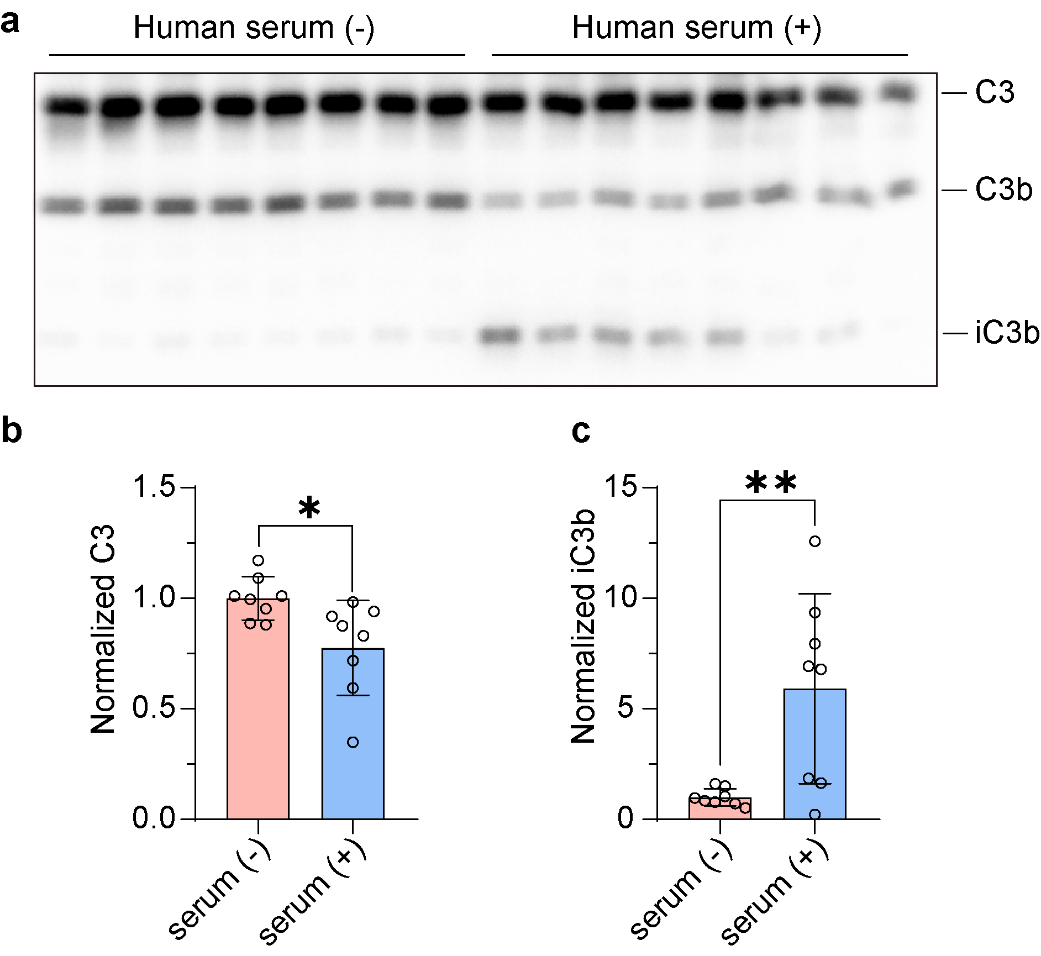

**Supplemental Figure 16. The cleavage of C3 protein in human serum. (a)** Western blot detection of C3 protein in untreated human serum. The quantification of C3 **(b)** and iC3b **(c)** by normalizing gray values. Data are means±SDs (n=8). Statistical difference is analyzed by Student's t-test with GraphPad Prism 9.0 (*^∗^p<0.05*, *^∗∗^p<0.01*).

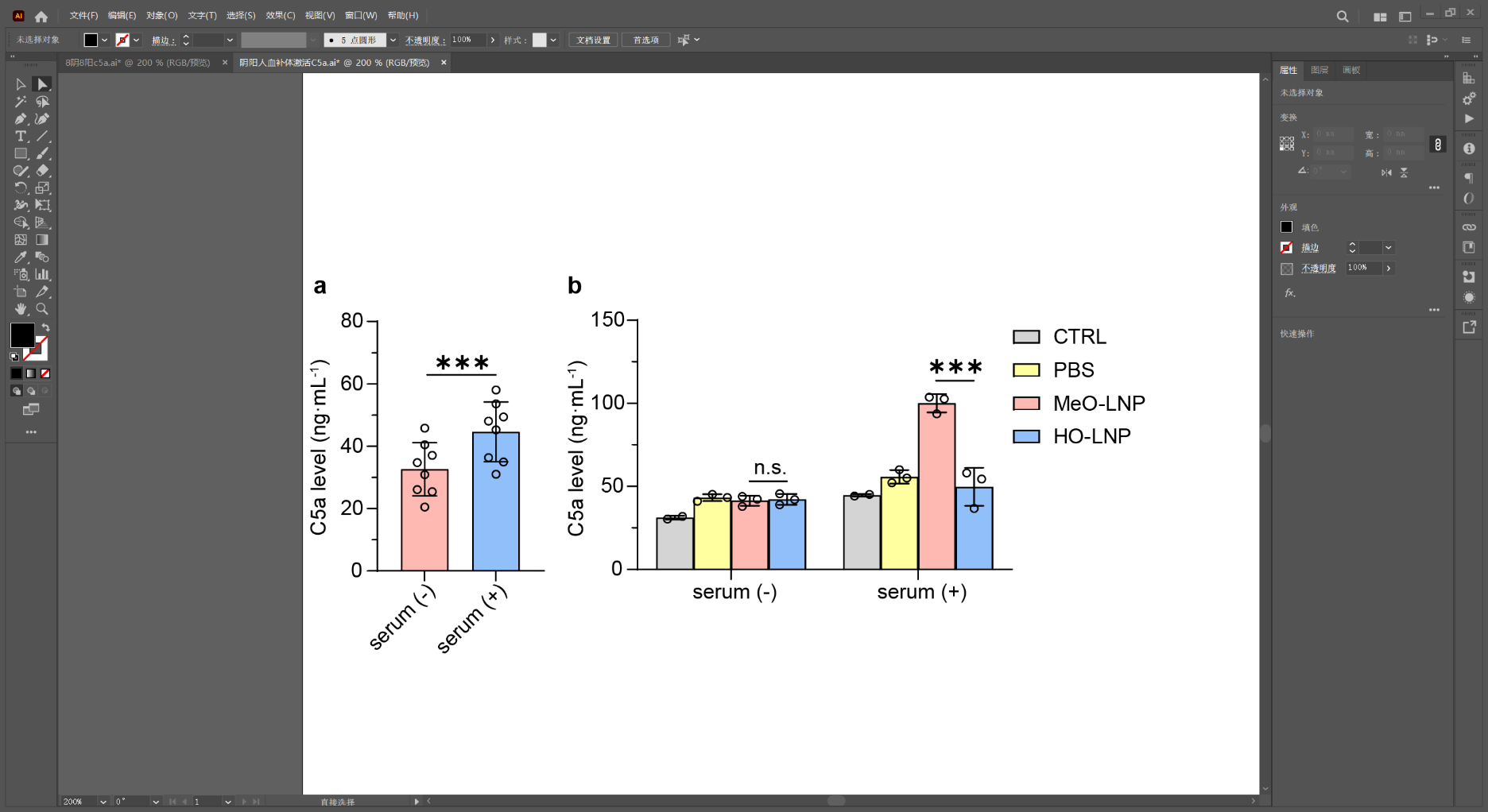

**Supplemental Figure 17. C5a level in the untreated human serum.** Data are means ± SDs (n = 8). Statistical difference is analyzed by Student's t-test with GraphPad Prism 9.0 (*^***^p<0.001*).

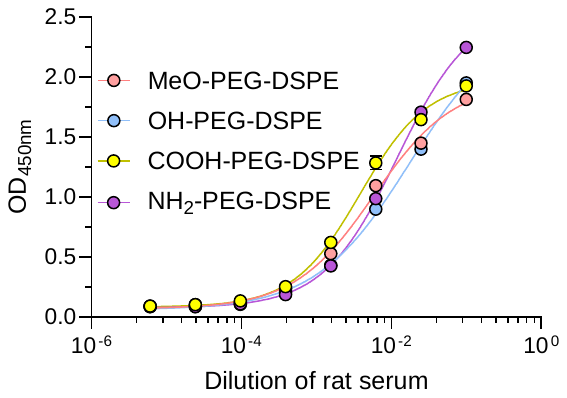

**Supplemental Figure 18. The binding of pre-stimulated anti-PEG IgM in rats towards PEG with different terminal groups.** SD rats were intravenously injected with empty PEGylated liposomes (2.5 mg HSPC/kg) to generate anti-PEG antibodies. After 6 days, blood was sampled and the serum was separated. The binding affinities of anti-PEG IgM towards PEG with different terminal groups were measured by sandwich ELISA assay (see Methods).

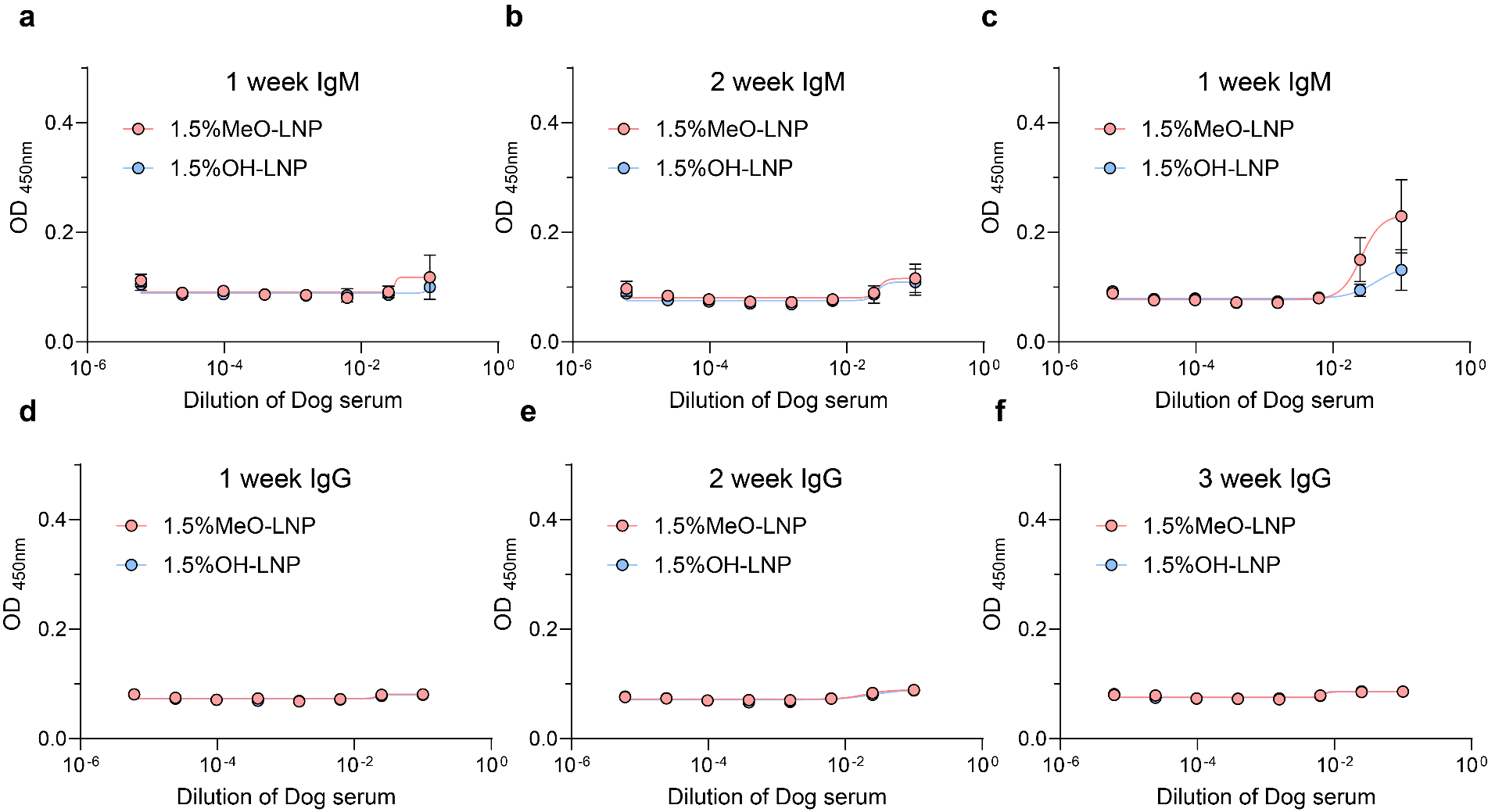

**Supplemental Figure 19. The immunogenicity of LNP in mice. (a-f)** The anti-PEG IgM and IgG was detected in dog serum from 1 week to 3 weeks after intravenous injection of MeO-LNP or OH-LNP weekly. Data are means±SDs (n=6) and analyzed with GraphPad Prism 9.0.

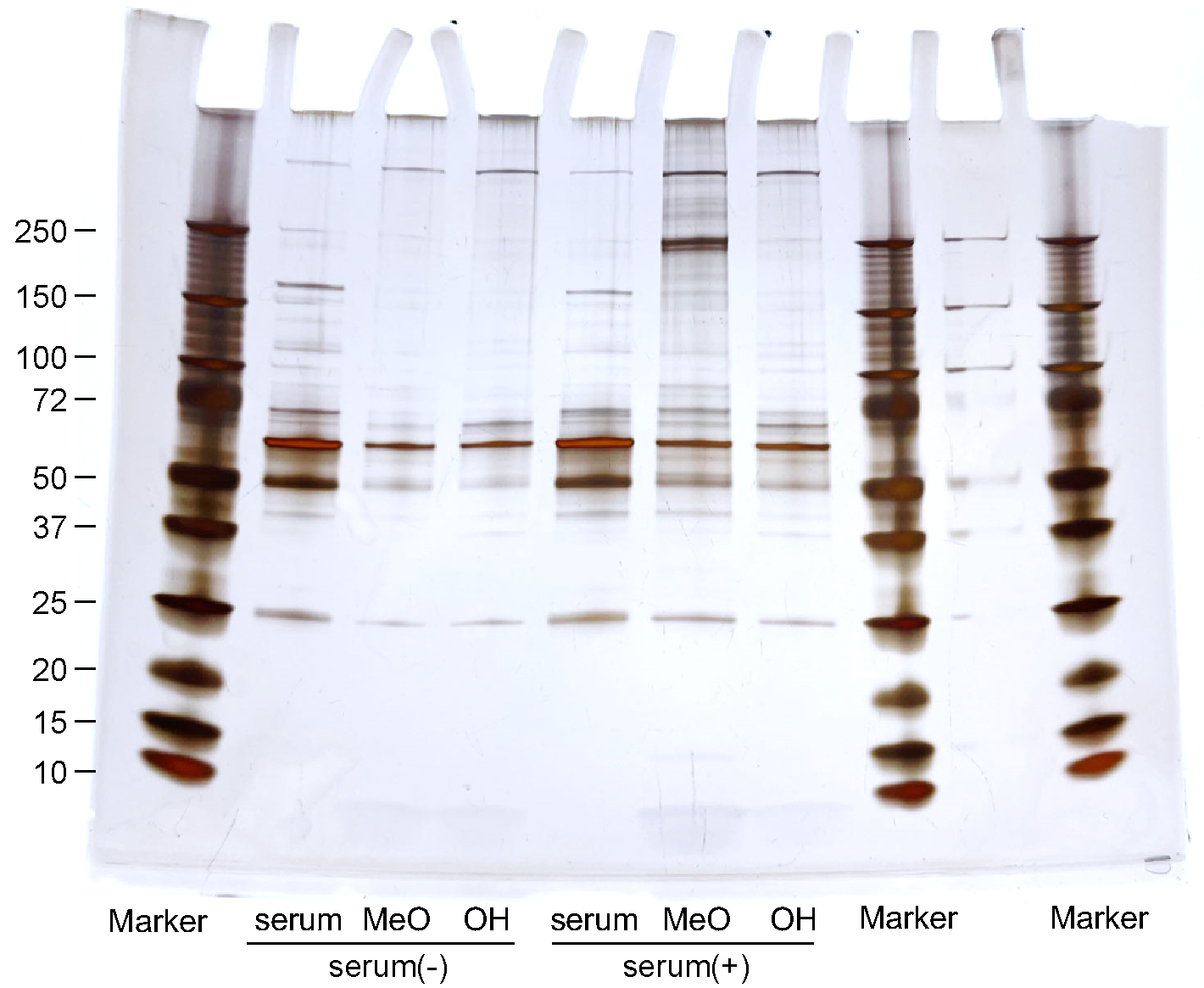

**Supplemental Figure 20. The uncropped and unprocessed gel with all size markers of Figure 4d.**

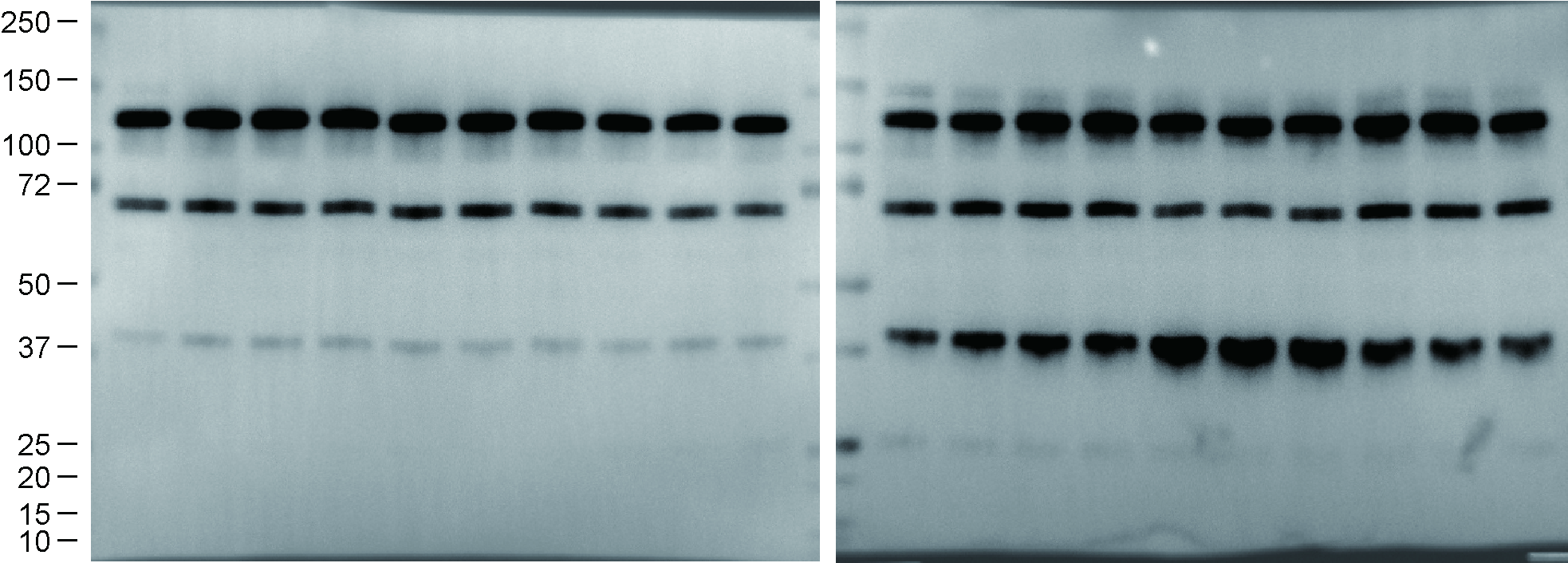

**Supplemental Figure 21. The uncropped and unprocessed gel with all size markers of Figure 5a and b.**
